# Stage-specific regulation of the *Plasmodium falciparum* proteasome activity reveals adaptive rewiring in artemisinin resistance

**DOI:** 10.1101/2025.11.21.689772

**Authors:** Shubha Bevkal Subramanyaswamy, Gourab Paul, Deborah Ogunribido, Andrea Mirauti, Daqiang Li, Wenhu Zhan, Gang Lin, Laura Kirkman

**Affiliations:** Division of Infectious Diseases, Department of Medicine, Weill Cornell Medicine, 1300 York Avenue, NY, 10065; Department of Microbiology and Immunology, Weill Cornell Medicine, 1300 York Avenue, NY, 10065

## Abstract

The ubiquitin-proteasome system (UPS) is essential for *Plasmodium falciparum* to maintain protein homeostasis, adapt to proteotoxic stress, and regulate parasite growth and stage transitions. The proteolytic 20S proteasome core is the central component of the UPS, where unfolded protein substrates are degraded into oligopeptides. Mechanisms regulating malaria parasite proteasome activity are poorly understood and have not been thoroughly studied. This knowledge gap is especially critical in the context of artemisinin (ART) resistance, where parasite survival depends on an enhanced stress response, including a greater reliance on the UPS. Here, we profiled proteasome activity and abundance across the parasite intraerythrocytic developmental cycle (IDC) in both ART-sensitive (ART-S) Dd2 and ART-resistant (ART-R) Dd2K13^R539T^ parasites. We uncovered striking stage-specific regulation: proteasome activity was abundant in the ring stage, decreased in trophozoites, and then peaked in schizonts. Furthermore, ART-R Dd2K13^R539T^ parasites exhibited higher ring-stage proteasome activity than ART-S Dd2, despite reduced proteasome abundance, suggesting a unique adaptive rewiring of proteasome function. To study proteasome regulation in the parasite, we manipulated proteasome abundance in Dd2 and Dd2K13^R539T^ by creating a conditional knockdown of PfUMP1, a conserved proteasome maturation factor. PfUMP1 depletion disrupted 20S assembly, decreased proteasome activity, and led to parasite death. These experiments uncovered two key features of proteasome regulation in *P. falciparum*: (1) the absence of canonical transcriptional regulation of proteasome genes in response to downregulation of proteasome activity, and (2) ART-R parasites exhibit a ring-stage specific increased sensitivity to proteasome downregulation. Together, our findings reveal a previously unrecognized layer of proteasome regulation in malaria parasites and how, as part of their survival adaptations to decreased hemoglobin uptake associated with ART resistance, these parasites alter proteasome function to survive. This work reinforces the therapeutic potential of the proteasome as a stage- and resistance-specific antimalarial target.

## Introduction

Malaria, caused by the apicomplexan parasite *Plasmodium falciparum (Pf),* remains a serious global health challenge^1^. Artemisinin combination therapy (ACT) is the current frontline treatment, and its potency has led to sharp decreases in malaria mortality. However, the emergence of artemisinin (ART) - resistant strains in Southeast Asia and, more recently and independently, in Africa threatens continued global efforts to combat malaria ^1–5^. Previous work demonstrated a critical role for the proteasome in response to the widespread cellular damage elicited by activated ART^6–8^. For this reason, we set out to provide a detailed study of the *Pf* proteasome across the intraerythrocytic development cycle (IDC) in otherwise isogenic ART-resistant (ART-R) and ART-sensitive (ART-S) lab strains^9^.

ART exerts its antimalarial effect after activation by heme, a byproduct of the degradation of host red blood cell hemoglobin, which is endocytosed by the parasite. Activated ART generates reactive oxygen species (ROS), causing widespread protein alkylation and cellular damage to biomolecules, leading to ER stress, inhibition of protein synthesis, accumulation of misfolded and/or polyubiquitinated proteins overwhelming UPS capacity, and ultimately, parasite death ^8,10–12^. ART resistance is predominantly associated with nonsynonymous mutations in the *P. falciparum kelch13* (PfK13) propeller region, resulting in decreased hemoglobin endocytosis and reduced ART^13–15^ activation. ART resistance is stage-specific and mostly restricted to the early ring stage of the intra-erythrocytic development cycle (IDC)^15^. Transcriptomic studies suggest that ART-R parasites achieve this by upregulating key components of the ubiquitin-proteasome system (UPS) and the unfolded protein response (UPR), thereby enhancing survival in the face of oxidative stress and protein damage^6,7,16^.

The importance of the UPS and the proteasome, particularly in response to ART-induced oxidative damage, has spurred the development of parasite-selective proteasome inhibitors as promising antimalarial candidates^17–20^. The proteasome is a cylindrical, multi-subunit complex composed of a 20S core particle (CP) and regulatory cap. The 20S CP, built of four stacked rings of seven α and β subunits with α_1-7_β_1-7_β_1-7_α_1-7_ organization, contains the catalytic subunits β1, β2, and β5, responsible for caspase, trypsin, and chymotrypsin-like activities, respectively^21^. Regulatory caps, such as the 19S, proteasome activator 28 (PA28), and proteasome activator 200 (PA200), modulate the access of ubiquitinated substrates to the catalytic core, all of which are conserved in *P. falciparum* ^22,23^. Damaged and/or ubiquitin-tagged proteins are routed to the proteasome, where they are unfolded and threaded into the 20S catalytic core for degradation. In this manner, the proteasome plays a key role in regulating cell development and can orchestrate and increase protein degradation capacity in response to cellular stress^23^.

Eukaryotes employ various mechanisms to modify proteasome abundance, composition, and activity. These include the dynamic remodeling of proteasome complexes, such as through association with alternative caps (19S/PA28/PA200), changes in cellular localization, transcriptional control of proteasome subunit expression by stress-responsive transcription factors like Rpn4 and Nrf2^24–29^, and modulation of proteasome function through diverse post-translational modifications. Proteasome regulation also occurs at the level of biogenesis, a highly coordinated process requiring dedicated chaperones. Among these, UMP1 plays a critical role in β-subunit maturation and in the assembly of the 20S CP^30,31^. Yeast cells lacking Ump1 exhibit defective coordination between β-subunit processing and later stages of proteasome assembly, resulting in functionally impaired proteasomes ^32^. In mammalian cells, UMP1 is essential ^31^.

Despite the importance of the proteasome to parasite survival and ART resistance, fundamental aspects of proteasome biology in *P. falciparum* remain unresolved. Previous work has mainly focused on the proteasome in response to ART treatment ^8,10,18,33,34^. Here, we provide the first systematic analysis of proteasome activity and abundance across the IDC in isogenic ART-Sand ART-R *P. falciparum*. We demonstrate that proteasome activity and abundance follow a dynamic, stage-specific cycle. By conditionally depleting PfUMP1, we establish its essential role in proteasome biogenesis and uncover two key insights: *P. falciparum* lacks canonical transcriptional compensation upon proteasome disruption, and ART-R parasites are more vulnerable to proteasome perturbation at the ring stage of the IDC.

## RESULTS

### Proteasome activity varies across the malaria parasite intraerythrocytic development cycle

We performed a direct characterization of proteasome activity across the IDC (Fig. 1A) at three distinct stages: rings (12–16 hours post-invasion, hpi), trophozoites (27–30 hpi), and schizonts (39–42 hpi) in ART-S Dd2 parasites using native gel electrophoresis with the activity-based substrate, Suc-LLVY-AMC, that detects ß5 chymotrypsin-like activity, to visualize distinct proteasome isoforms (30S, 26S and 20S) in all three stages of the IDC (Fig. 1B)^35^. Equal amounts of protein from the three parasite development stages were loaded to reflect proteasome activity per total protein in a parasite lysate. Additionally, we treated the gel with SDS to assess the presence of uncapped 20S isoforms, which are otherwise kept in a closed state as the outer alpha rings prevent the probe from entering the 20S core. This revealed a strong band and the presence of the 20S core at all three stages of the IDC (Fig. 1B). Proteasome activity peaked in the schizont stage, with both dually capped (30S) and single-capped (26S) proteasomes contributing to overall activity, paralleling published evidence that protein ubiquitination peaks in schizonts^36^. Notably, substantial proteasome activity was also detected at the ring stage, with the lowest proteasome activity in the trophozoite stage. For comparison, if we consider capped proteasome activity (30S and 26S only, not including free 20S) in schizonts as 100%, rings and trophozoites showed 47% (±5% SD) and 27% (±4.7% SD), respectively.

**Figure 1:**
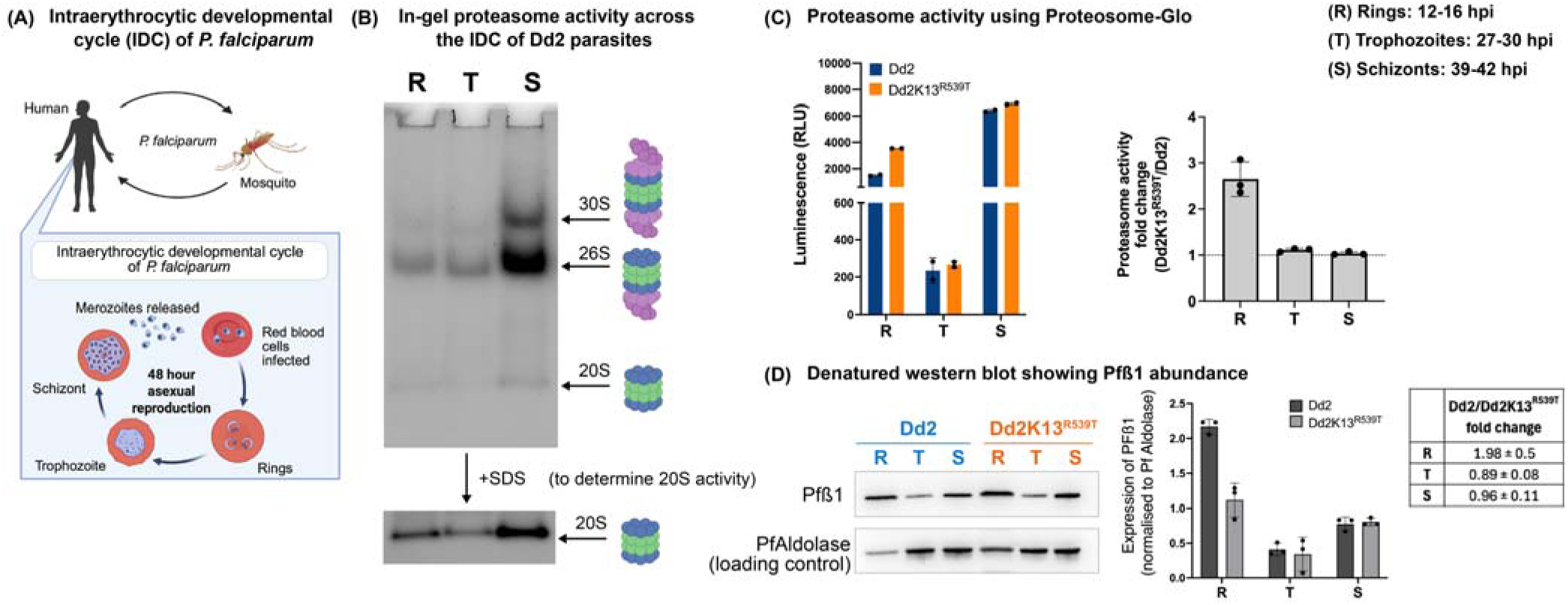
Proteasome activity and expression across the *P. falciparum* IDC. **(A)** A Schematic of intraerythrocytic development cycle (IDC) of *P. falciparum.* Parasite shuttles between mosquito and human host. *P. falciparum* infection starts when an infected Anopheles mosquito injects sporozoites into the human bloodstream. These migrate to the liver, multiply, and release merozoites that invade red blood cells. Within red blood cells, parasites progress from ring to trophozoite to schizont stages, releasing new merozoites upon cell rupture to continue the IDC. Some parasites develop into gametocytes, which are taken up by mosquitoes, completing the sexual phase of the life cycle. **(B)** In-gel proteasome activity assay using SucLLVY-AMC (detects chymotrypsin activity exhibited by the ß5 subunit of the proteasome) as substrate. Extracts from Dd2 parasites (rings, trophozoites and schizonts) were separated by native gel and peptidase activity from the proteasome was visualized using SucLLVY-AMC in gel activity assay in the absence (top) or presence (bottom) of 0.02% SDS (which opens and activates the 20S proteasome to visualize free 20S core particle). Gel is a representative of three biological replicates (see **Supplemental Fig. 1A** for biological replicates). **(C)** Proteasome-Glo^TM^ assay on rings (12-16 hpi), trophozoites (27-30 hpi) and schizont (39-42 hpi) stages of Dd2 and Dd2K13^R539T^ parasites. Cellular lysates were incubated with Suc-LLVY-AMC for 10 min at 37°C, and fluorescence intensity was measured. Graph is a representative of three biological replicates (see **Supplemental Fig. 1B** for biological replicates). Right panel: Proteasome activity fold change (Dd2K13^R539T^/Dd2) in rings, trophozoites and schizonts from 3 biological replicates. **(D)** Western blot analysis of Pf proteasome ß1 subunit expression levels across the IDC (rings (12-16 hpi), trophozoites (27-30 hpi) and schizont (39-42hpi) stages) of Dd2 and Dd2K13^R539T^ parasites. PfAldolase served as loading control. Blot is a representative of three independent biological replicates. Right panel: western blot quantification of Pf proteasome ß1 subunit expression from three biological replicates (see **Supplemental Fig.1C** for biological replicates).

### ART-R ring stage parasites exhibit elevated proteasome activity despite reduced subunit abundance

Given that ART resistance in *P. falciparum* is closely associated with changes in proteasome activity, particularly in the parasite’s response to oxidative and proteotoxic stress ^6,16,34^, we examined proteasome activity in ART-R Dd2K13^R539T^ parasites, which differ from ART-S Dd2 only by the PfK13^R539T^ mutation conferring ART resistance. Native gel-based proteasome activity assays requires large amounts of protein (∼100µg), so to overcome this limitation, we employed the sensitive luminescent Proteasome-Glo assay (Promega), which detects chymotrypsin-like activity across the 30S, 26S, and 20S proteasome isoforms from as little as one µg lysate^37^. Synchronous cultures of Dd2 and Dd2K13^R539T^ parasites were harvested at ring, trophozoite, and schizont stages of the IDC described above. As shown in Fig. 1C, Dd2K13^R539T^ rings exhibited significantly elevated proteasome activity (2.6-fold change, ±0.37 SD) relative to Dd2 rings. At the same time, no significant differences were observed at trophozoite (1.1-fold change, ±0.03 SD) or schizont stages (1.04-fold change, ±0.03 SD). These results, obtained under standard culture conditions, represent the baseline proteasome activity profile of both parasite lines across the IDC.

To determine whether altered proteasome activity correlated with proteasome abundance, we generated an antibody against the Pfβ1 subunit and quantified Pfβ1 protein levels at the same developmental stages (Fig. 1D). With equal protein loading and normalization to the housekeeping protein PfAldolase, we found that in ART-S Dd2 parasites, Pfβ1 abundance was significantly higher in rings (5.4-fold change, ±0.9 SD) and schizonts (2-fold change, ±0.3 SD) compared to trophozoites, consistent with the reduced activity observed in trophozoites. We then confirmed this pattern was consistent across different parasite lines by repeating this work using the 3D7 laboratory strain (Supplementary Fig. 2A).

When directly comparing ART-S Dd2 and ART-R Dd2K13^R539T^ parasites, we observed no differences in Pfß1 abundance at trophozoite (0.9-fold change, ±0.08 SD) and schizont stages (0.96-fold change, ±0.11 SD) (Fig. 1D). In contrast, in the ring stage, Pfβ1 levels were consistently lower in ART-R parasites (2-fold change, ±0.55 SD) than in ART-S parasites (Fig. 1D). This finding was reproducible across three biological replicates and was validated using an independent antibody recognizing 20S core α-subunits (Supplementary Fig. 2B & C).

It is important to note that these measurements reflect subunit abundance under basal culture conditions and do not directly correspond to the abundance of fully assembled, catalytically active proteasomes. However, among the seven β-subunits of the proteasome, β1, β2, and β5–β7 undergo N-terminal processing to yield mature, active subunits incorporated into the proteasome core. The in-house–generated anti-Pfβ1 antibody specifically recognized the mature, processed β1 subunit in rings and trophozoites, with a faint band corresponding to unprocessed β1 detected in schizonts of both ART-S Dd2 and ART-R Dd2K13^R539T^ parasites. Thus, our quantification of proteasome abundance across the IDC reflects levels of the processed, mature Pfβ1 subunit s(Fig. 1D).

Together, these results reveal a paradox in which ART-R ring stage parasites exhibit elevated proteasome activity despite reduced proteasome subunit abundance. This, in turn, suggests that additional regulatory mechanisms beyond subunit quantity modulate proteasome activity in ART-R parasites and highlights the importance of altered proteasome biology in the absence of mutations in 20S core or 19S cap subunits.

### Pre-assembled proteasomes are present in merozoites and essential for *P. falciparum* ring-stage development

Mature trophozoites undergo 3–6 rounds of nuclear and cytoplasmic division to form schizonts containing ∼8–36 merozoites^38,39^. When mature schizonts rupture, the released merozoites invade fresh erythrocytes and initiate the next erythrocytic cycle (Fig. 1A). We hypothesized that proteasome assembly occurs during the trophozoite-to-schizont transition and that newly formed merozoites are pre-loaded with assembled proteasomes, which are subsequently inherited by ring-stage parasites, accounting for ring stage proteasome activity seen above. To test this, we isolated tightly synchronized 44–46 hpi segmented schizonts of ART-S Dd2 parasites and treated them with 2.5 μM compound 1 for 5 hours. Compound 1 blocks schizont rupture without impairing merozoite formation, and this effect is reversible upon drug washout ^40^. Following compound 1 treatment, parasites were exposed to the covalent proteasome inhibitor WLW-VS (1 μM, 1 h) to irreversibly inhibit assembled proteasomes ^20^. Controls included DMSO and the reversible proteasome inhibitor TDI-8304 (1 μM) ^33^. Cultures were then washed free of inhibitors, returned to growth, and development was monitored for 3 days (Fig. 2A).

**Figure 2:**
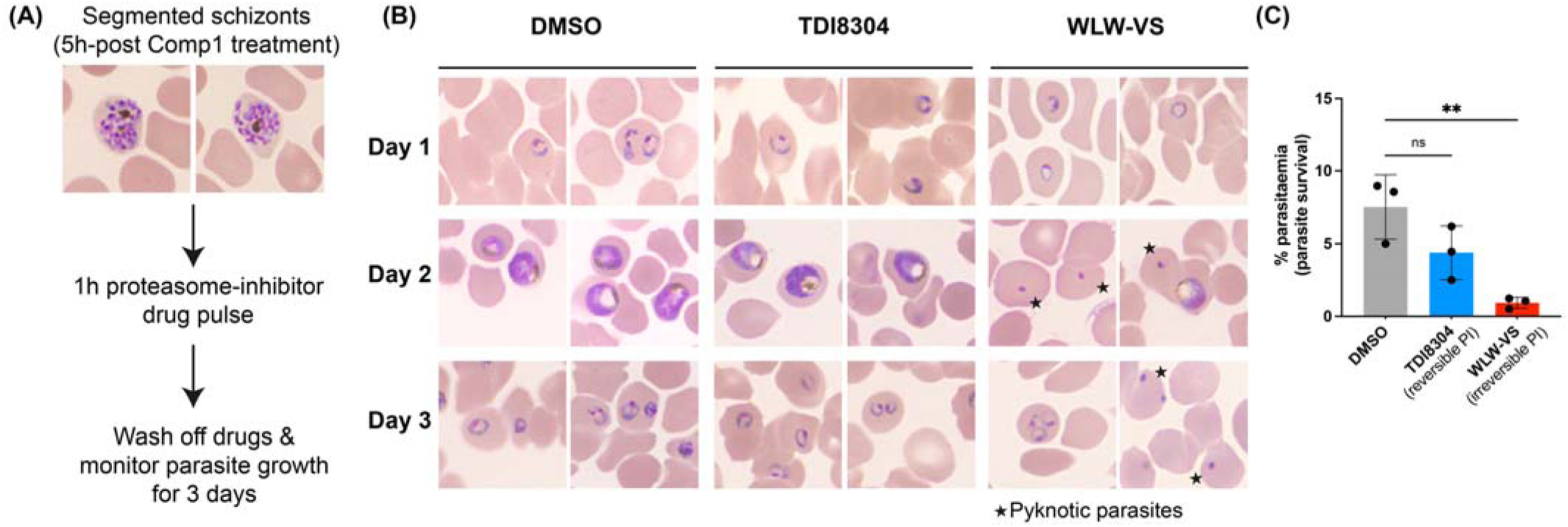
Ring-stage parasite viability depends on proteasomes assembled prior to egress. **(A)** Schematic of the experiment conducted. Briefly, tightly synchronized 44–46 hpi segmented schizonts of ART-S Dd2 parasites were treated with 2.5 μM compound 1 for 5 hours. Following compound 1 treatment, parasites were exposed to the covalent proteasome inhibitor WLW-VS (1 μM, 1 h) to irreversibly inhibit assembled proteasomes. Controls included DMSO and the reversible proteasome inhibitor TDI-8304 (1 μM). Cultures were then washed free of inhibitors and returned to growth, and development was monitored for 3 days. **(B)** Giemsa -stained smears of DMSO/TDI8304/WLW-VS treated parasites on days 1-3 post inhibitor treatment (as described in (A)). Stars represent pyknotic parasites. **(C)** Parasite survival on day 3 post of DMSO/TDI8304/WLW-VS treatment. Dots represent data from 3 biological replicates (See **Supplemental** Fig.3 for biological replicates)

DMSO-treated schizonts and those exposed to the reversible inhibitor TDI-8304 successfully egressed, released infective merozoites, and formed ring-stage parasites by day 1 (Fig. 2B). Parasites progressed normally through the IDC, producing trophozoites and schizonts on day 2 and new rings on day 3 (Fig. 2B). In contrast, WLW-VS–treated schizonts egressed and formed rings by day 1 but failed to develop further. By days 2 and 3, most parasites appeared pyknotic, with very few viable trophozoites or rings (Fig. 2B). Quantification at day 3 confirmed a significant reduction in parasitemia in WLW-VS–treated cultures relative to DMSO controls (Fig. 2C). Together, these results support a model in which proteasome assembly prior to merozoite egress primes early rings with the proteolytic capacity required to withstand intracellular stress and initiate development.

### PfUMP1 is a proteasome assembly factor and essential for parasite survival

To manipulate proteasome activity in the parasite without using inhibitors, we used an inducible knockdown system targeting the proteasome maturation factor, PfUMP1 (PfDd2_050032900), to modulate proteasome levels and assess its impact on parasite development in the ART-S Dd2 line. pSLI-HA-T2A-NeoR-glmS selection-linked integration method was used to integrate HA and glmS ribozyme sequence to the 3’ end of the Pf UMP1 gene (Fig. 3A) and integration was confirmed by PCR (Supplementary Fig. 4)^41^. In this transgenic line, PfUMP1 mRNA expression is under the control of the glmS ribozyme, which induces transcript degradation in the presence of glucosamine (GlcN) (Fig. 3B). The insertion of HA-T2A-NeoR-glmS sequence to the PfUMP1 gene resulted similar though slightly delayed parasite growth (growth rate: 2.19 ±0.08 per 48h IDC) compared to the parental Dd2 control (growth rate: 2.46 ± 0.16 per 48h IDC) (Fig. 3C)^41^. To assess the effect of GlcN on PfUMP1 protein levels, synchronized ring-stage cultures were treated with 0.5 mM GlcN for 24 hours.

**Figure 3:**
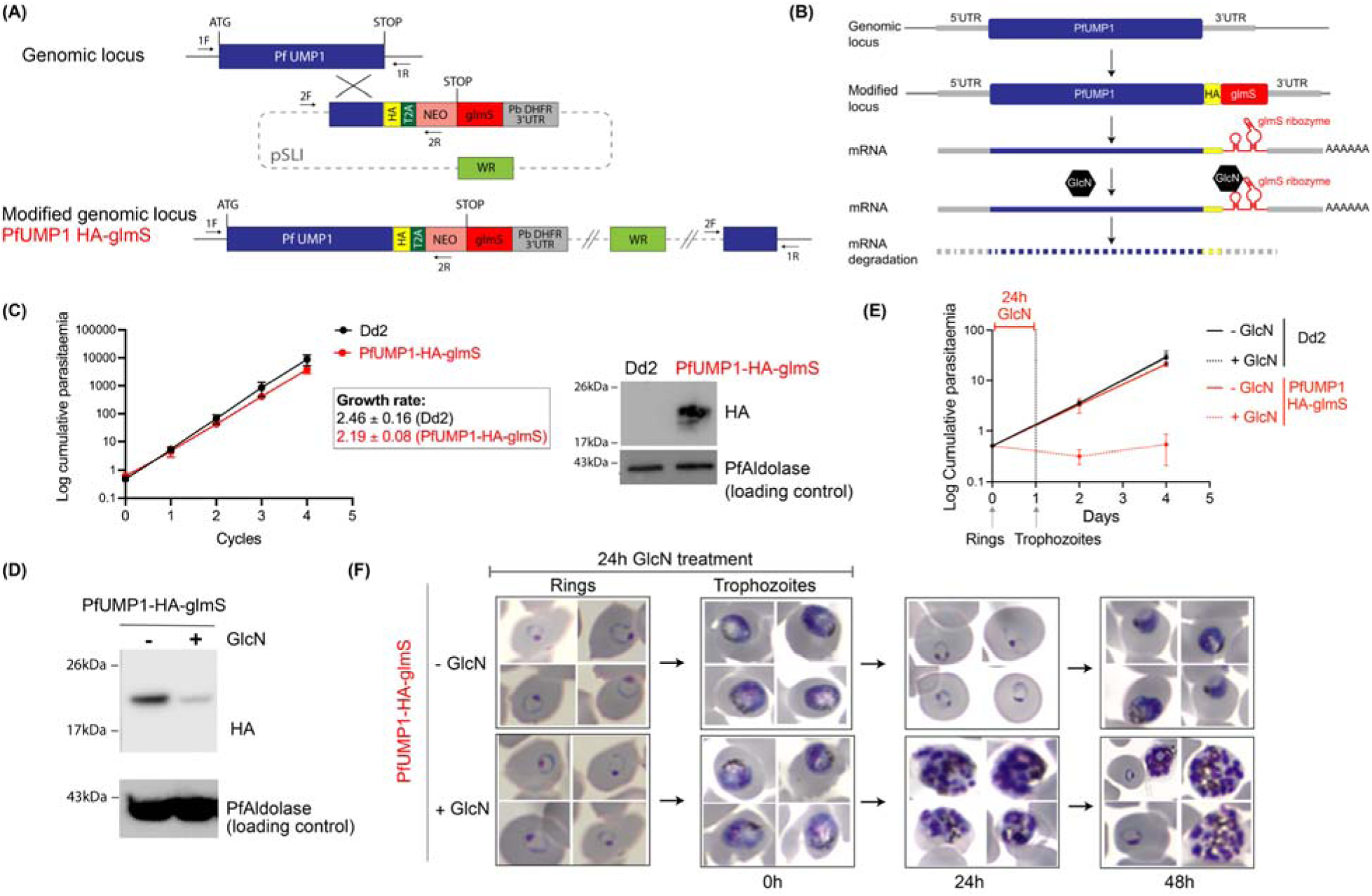
Conditional knockdown of PfUMP1 impairs parasite growth and development. **(A)** Schematic of the selection-linked integration-based strategy used to generate PfUMP1-HA-glmS parasites by single-crossover recombination. Arrows denote diagnostic PCR primers. T2A, skip peptide; Neo, neomycin-resistance gene; glmS, glmS ribozyme. **(B)** Schematic of the glmS ribozyme reverse genetic tool. The glmS ribozyme is inserted in the 3′-UTR after the coding region so that it is present in the expressed mRNA. Following addition of the inducer, glucosamine (GlcN), which binds to the ribozyme, the mRNA self-cleaves resulting in degradation of the mRNA and knock down of protein expression. **(C)** Cumulative growth curves of Dd2 and PfUMP1-HA-glmS parasites (n=3, mean ± SD). Inset: Western blot showing PfUMP1 expression detected with anti-HA antibody in PfUMP1-HA-glmS parasites, with Dd2 as a negative control. PfAldolase served as loading control. **(D)** Western blot analysis showing ∼80% reduction in PfUMP1 protein levels after treatment with 0.5 mM GlcN for 24 h at the ring stage in PfUMP1-HA-glmS parasites. PfAldolase served as loading control. **(E)** Cumulative growth analysis of PfUMP1-HA-glmS parasites with or without a 24 h GlcN pulse to induce PfUMP1 knockdown. Dd2 served as a negative control (n=3, mean ± SD). **(F)** Giemsa-stained smears of PfUMP1-HA-glmS parasites following a 24 h GlcN pulse to induce knockdown, followed by GlcN washout. Upper panel: uninduced parasites; lower panel: knockdown-induced parasites.

Western blot analysis showed an 82% reduction in PfUMP1 levels in the GlcN-treated PfUMP1-HA-glmS parasites, indicating efficient knockdown (Fig. 3D). Next, we evaluated the impact of PfUMP1 knockdown on parasite growth by treating synchronized Pf-UMP1-HA-glmS ring-stage parasites with or without 0.5 mM GlcN for 24 hours, then removing GlcN and monitoring growth over four days. GlcN treatment significantly impaired parasite growth, as quantified by flow cytometry, indicating that PfUMP1 is essential for parasite survival (Fig. 3E). Examination of Giemsa-stained smears showed that while the GlcN-treated PfUMP1-HA-glmS parasites developed normally through the trophozoite stage, then arrested in schizont stages compared to untreated control cultures that completed the IDC and reinvaded to form ring-stage parasites (Fig. 3F). Although some GlcN-treated parasites formed new rings by day 4, a substantial number remained arrested as schizonts as indicated by lack of growth seen in Fig. 3E. These findings are consistent with previous studies showing that disruption of the UPS impairs schizont maturation and merozoite invasion^36^.

Having established the efficiency of PfUMP1-glmS knockdown and the observed growth phenotype, we examined proteasome activity at the early schizont stage (34-39 hpi) of Dd2 and PfUMP1-HA-glmS parasites with or without GlcN using native gel electrophoresis described above. We found that the untreated PfUMP1-HA-glmS parasites showed ∼ 80 % proteasome activity compared to Dd2 and GlcN-treated PfUMP1-HA-glmS parasites displayed over a 95% reduction in proteasome activity compared to Dd2 (Fig. 4A). As a control, we treated Dd2 and PfUMP1-HA-glmS parasites with TDI-8304, a selective Plasmodium proteasome inhibitor ^33^, TDI-8304 reduced proteasome activity by more than 90% (Fig. 4A). These findings confirm that PfUMP1 regulates proteasome abundance and activity in *P. falciparum* and reduced proteasome interferes with the parasite’s ability to complete the IDC.

**Figure 4:**
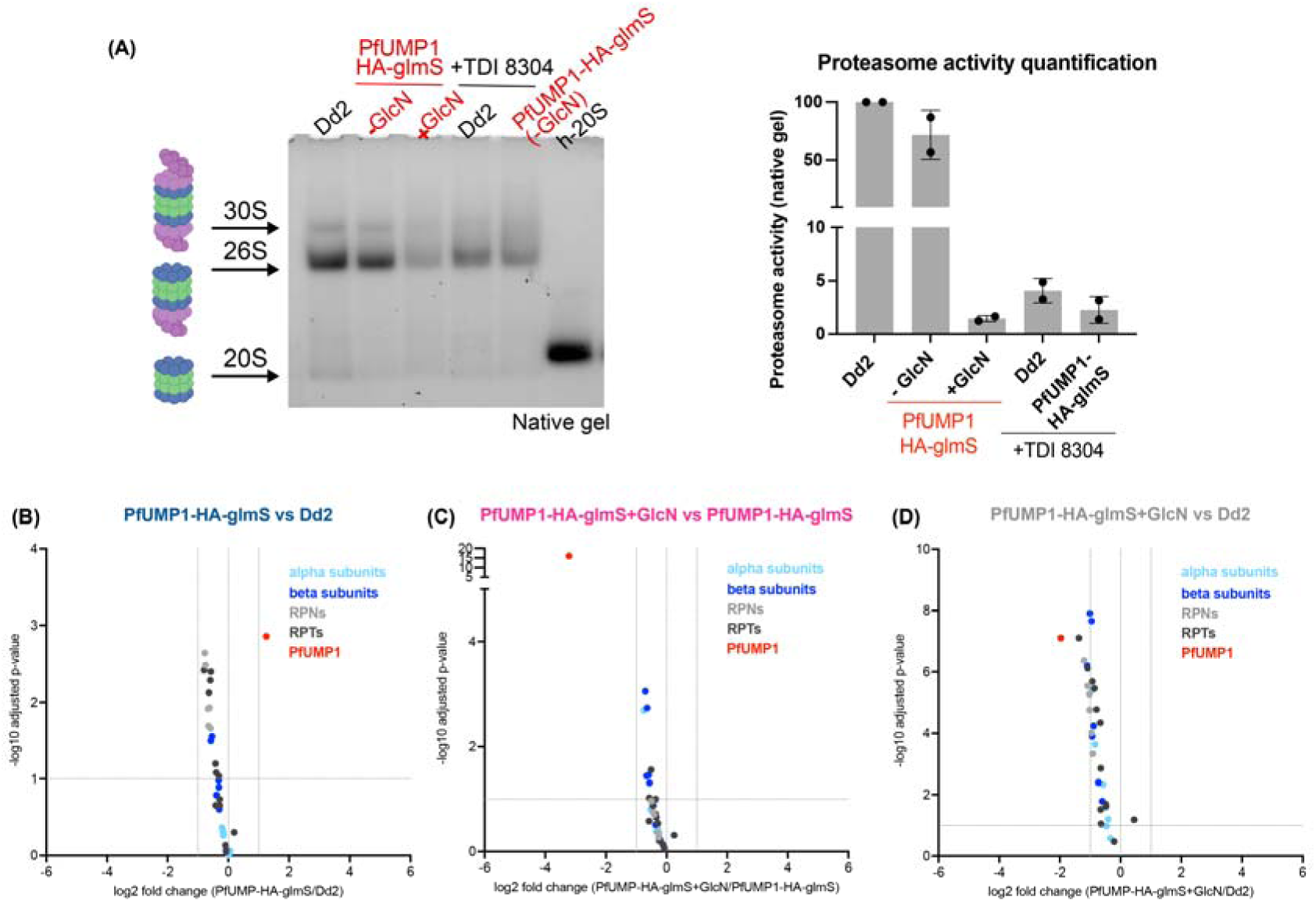
PfUMP1 knockdown impairs proteasome activity without altering proteasome transcript levels. **(A)** In-gel proteasome activity assay using SucLLVY-AMC (detects chymotrypsin activity exhibited by the ß5 subunit of the proteasome) as substrate. Cell extracts from 34-39 hpi Dd2 and PfUMP1-HA-glmS parasites, treated with or without GlcN for 24h, were separated by native gel and peptidase activity from the proteasome was visualized using SucLLVY-AMC. Analysis of proteasome activity of Dd2 and PfUMP1-HA-glmS parasites treated with TDI-8304, a selective plasmodium proteasome inhibitor served as control. Gel is a representative of two biological replicates (see **Supplemental** Fig.5 for biological replicates). Right: Quantitation of Native gels (data from **A**) **(B) – (D)** Transcriptome analysis upon PfUMP1 knockdown. Volcano plots comparing the Pf proteasome transcripts of Dd2, PfUMP1-HA-glmS (-GlcN) and Pf UMP1-HA-glmS (+GlcN). Data were obtained from 3 biological replicates. Vertical dotted lines indicate a fold change of ≥ 1 and horizontal dotted line indicates a p-value < 0.01. Dark blue dots, proteasome ß subunits; light blue dots, proteasome α subunits; light grey and dark grey subunits, proteasome cap subunits RPNs and RPTs; red dot, PfUMP1.

### *P. falciparum* lacks a canonical transcriptional response when proteasome activity is compromised

Proteasome activity and abundance are tightly regulated and responsive to different cell-cycle states and stressors. In yeast, mammals, and plants, the controlled expression of proteasome subunit genes is orchestrated by distinct but functionally analogous transcription factors: Rpn4, NRF1/2, and NAC53/78, respectively ^42^. These transcription factors bind to specific regulatory sequences within the promoter regions of proteasome subunit genes, modulating their expression during development or in response to stress, particularly when proteasome capacity is impaired or exceeded ^26,29,43–45^. In *P. falciparum*, although a homolog of Rpn4 (PF3D7_0317800) has been annotated, its role remains unstudied, and no homologs of NRF1/2 or NAC53/78 have been identified to date. Sequence analysis of the upstream regions (1500 nucleotides upstream of the translational start site) of *P. falciparum* proteasome subunit genes revealed no canonical regulatory sequences such as PACE, ARE, or PRCE. Other work to identify unique regulatory elements in *P. falciparum* found that genes encoding proteasome subunits were co-expressed and identified three strong motifs in the upstream regions of *P. falcipaum* proteasome subunit genes: CACA, a G-rich motif, and TGTG ^46^. Large-scale transcriptomic data from multiple studies also demonstrated that proteasome subunit transcription is regulated across the parasite IDC, with a peak expression at 28-34 hpi as the parasite transitions from trophozoite to schizont (Supplementary Fig.6).

Given the importance of transcriptional regulation in proteasome activity in higher eukaryotes, we investigated whether there might be unique transcriptional regulation in *P. falciparum* proteasome subunits. To this end, we utilized the PfUMP1-HA-glmS transgenic parasite line to modulate proteasome activity and assess transcriptional responses to reduced proteasome function. Knockdown of PfUMP1 was achieved by treating PfUMP1-HA-glmS parasites with 0.5 mM GlcN, and whole transcriptome analysis was performed on GlcN-treated PfUMP1-HA-glmS parasites alongside control lines (Dd2 and untreated PfUMP1-HA-glmS parasites). Parasites were harvested at 31–34 hpi for RNA-seq analysis. This time point was selected to ensure that GlcN-treated PfUMP1-HA-glmS parasites, which exhibit slightly slower progression through the IDC even without GlcN, were at the same developmental stage as the control parasites (Fig. 3C & E), thereby allowing the effects of reduced proteasome activity to be distinguished from potential differences in life cycle progression.

RNA-seq analysis revealed no significant changes in the mRNA expression levels of proteasome subunits following PfUMP1 knockdown, except for the expected decrease in PfUMP1 transcript levels (Fig. 4B-D). Moreover, no significant changes in the expression of other UPS-related genes were observed. These findings suggest a lack of transcriptional regulation of proteasome subunit genes in *P. falciparum* under conditions of reduced proteasome activity and highlight the possibility that proteasome regulation in *P. falciparum* likely occurs via alternative mechanisms, such as post-translational modifications, rather than through canonical transcriptional control.

### ART-R parasites are highly sensitive to reduced proteasome activity

We next investigated the impact of reduced proteasome activity in ART-R parasites. For this, we again selected Dd2K13^R539T^ parasites. Using the same PfUMP1-HA-glmS knockdown strategy, we generated a conditional PfUMP1 knockdown line in the Dd2K13 ^R539T^ background. Under normal growth conditions (without GlcN), Dd2K13 ^R539T^ PfUMP1-HA-glmS parasites exhibited a pronounced growth defect (growth rate: 1.68 ± 0.10 per 48h IDC) compared to the parental Dd2K13 ^R539T^ control (growth rate: 2.39 ± 0.20 per 48h IDC), which was greater than that seen in the Dd2 versus Dd2 PfUMP1-HA-glmS lines (Fig. 5A). Addition of 0.5mM GlcN induced effective PfUMP1 knockdown (∼88% reduction, as quantified by western blot) and addition of GlcN revealed PfUMP1 to be essential for parasite growth in the ART-R parasites (Fig. 5B). Our data suggest a heightened sensitivity to reduction in assembled proteasomes in ART-R parasites.

**Figure 5:**
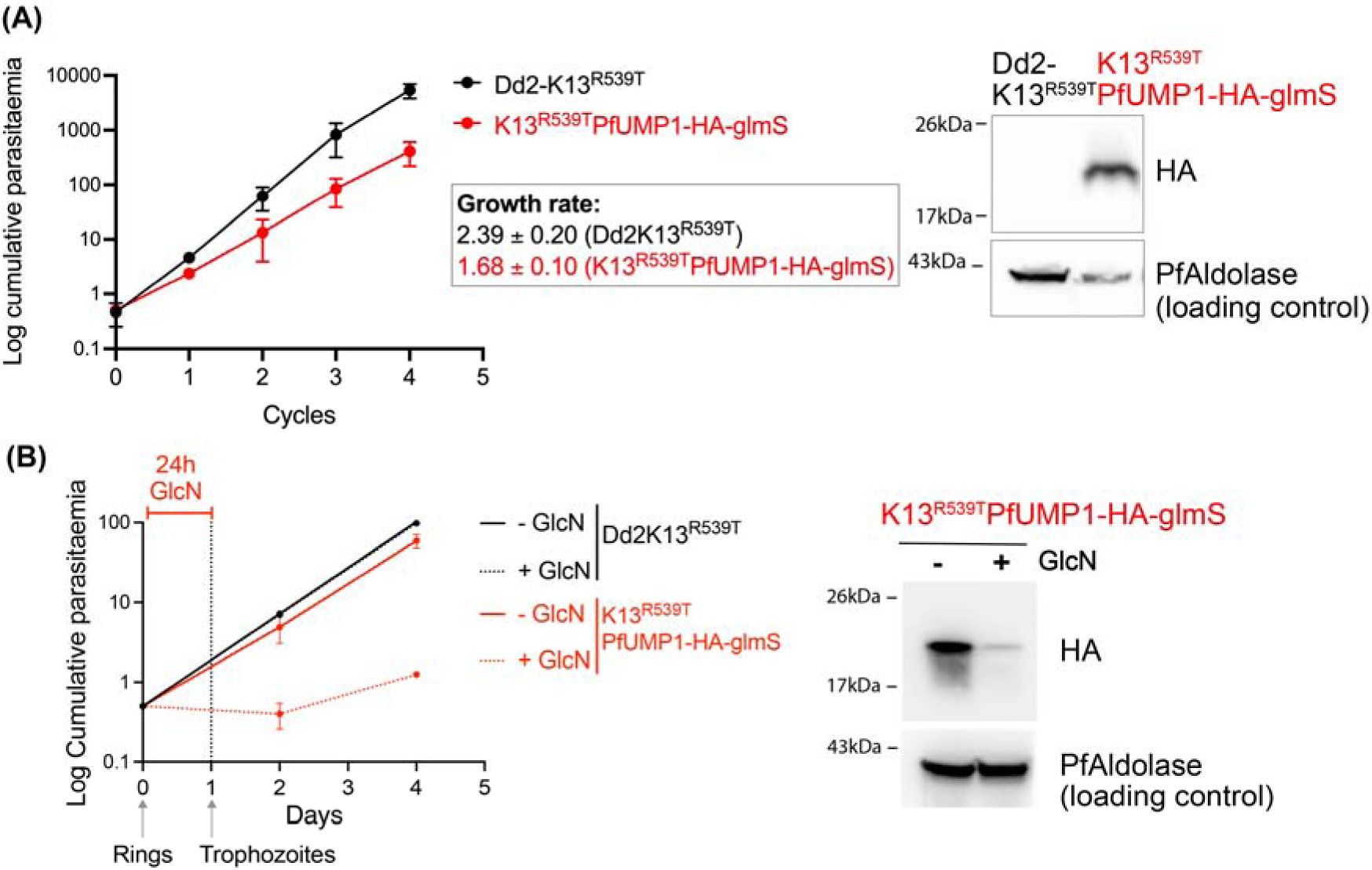
PfUMP1 is required for survival of ART-resistant parasites. **(A)** Cumulative growth curves of Dd2-K13^R539T^ and K13^R539T^PfUMP1-HA-glmS parasites (n=3, mean ± SD). Right panel: Western blot showing PfUMP1 expression detected with anti-HA antibody in K13^R539T^PfUMP1-HA-glmS parasites, with Dd2-K13^R539T^ as a negative control. PfAldolase served as loading control. **(B)** Cumulative growth analysis of K13^R539T^PfUMP1-HA-glmS parasites with or without a 24 h GlcN pulse to induce PfUMP1 knockdown. Dd2-K13^R539T^ served as a negative control (n=3, mean ± SD). Right panel: Western blot analysis showing reduction in PfUMP1 protein levels after treatment with 0.5 mM GlcN for 24 h at the ring stage in K13^R539T^PfUMP1-HA-glmS parasites. PfAldolase served as loading control.

### Parasite sensitivity to UMP1 knockdown varies across the IDC, with prolonged sensitivity in ART-R parasites

To evaluate the effect of variable proteasome activity during the parasite IDC, we used highly synchronized parasites and pulsed them with GlcN at four different time points, as shown in Fig. 6A. Since PfUMP1 is essential for parasite survival, we applied our previous method of pulsing cultures with GlcN for 24 hours and observing parasite recovery. PfUMP1 knockdown reduces the assembly of new 20S cores but does not affect the function of already assembled proteasomes.

**Figure 6:**
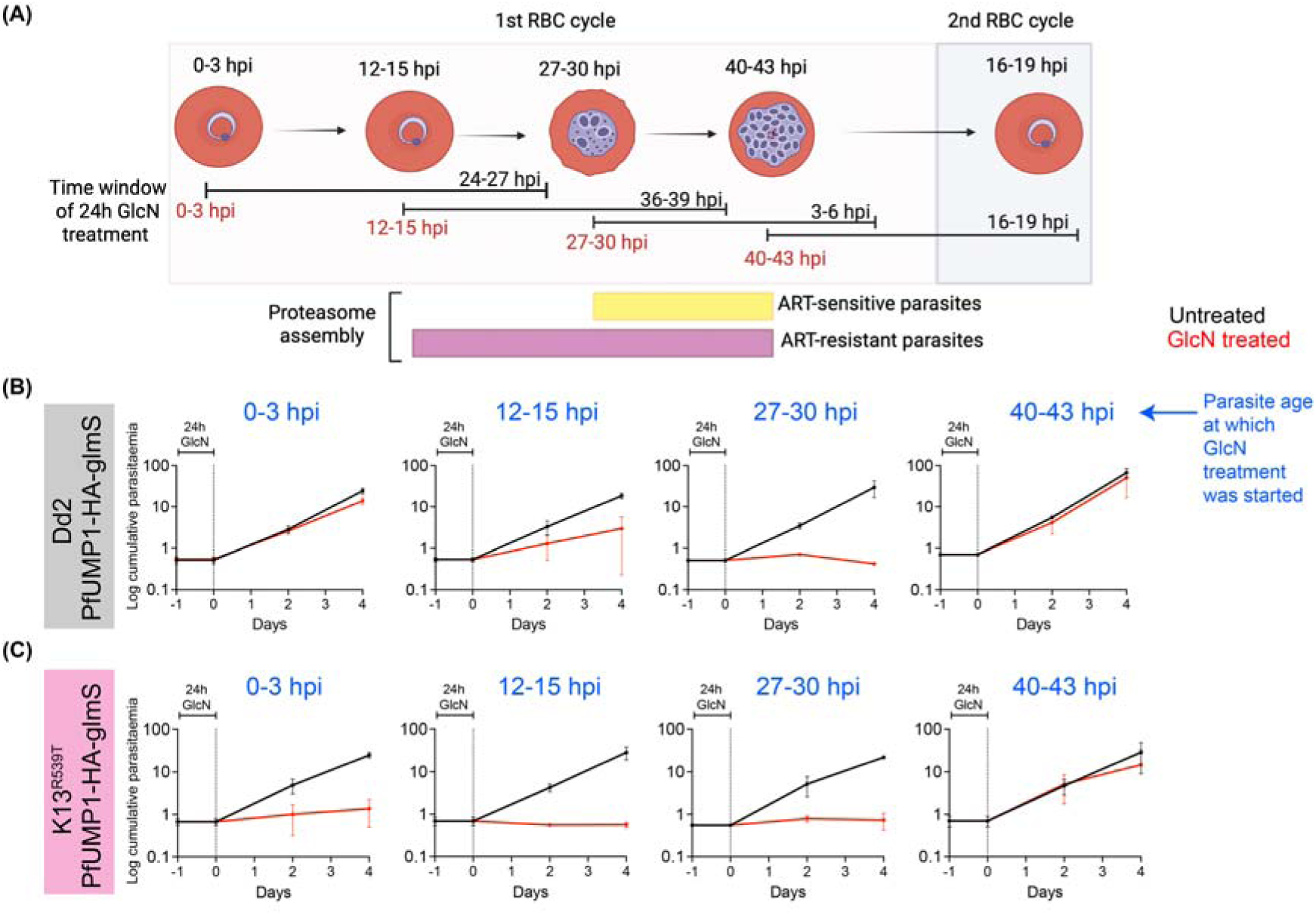
Stage-specific sensitivity to PfUMP1 depletion in Dd2 and ART-resistant parasites. **(A)** Schematic of PfUMP1 knockdown across the IDC. Dd2-PfUMP1-HA-glmS and K13^R539T^PfUMP1-HA-glmS cultures were treated with 0.5 mM GlcN or RPMI control for 24h at distinct developmental stages (0–3 hpi, 12–15 hpi, 27–30 hpi, and 40–43 hpi). GlcN was then washed out, and parasite growth was monitored by flow cytometry every 48 h for 4 days. **(B)** Cumulative growth analysis of Dd2-PfUMP1-HA-glmS parasites following a 24 h GlcN pulse at different IDC stages (n=3, mean ± SD). **(C)** Cumulative growth analysis of K13^R539T^PfUMP1-HA-glmS parasites following a 24 h GlcN pulse at different IDC stages (n=3, mean ± SD).

ART-S Dd2 parasites were most sensitive to PfUMP1 knockdown during the late ring to mid-trophozoite stages, when parasites experienced 24-hour GlcN treatment at 12-15 hpi and 27-30 hpi (Fig. 6A). In contrast, there was minimal impact on parasite survival when GlcN was added during the early ring (0-3 hpi) or schizont stages (40-43 hpi) (Fig. 6B). This indicates that critical proteasome assembly in ART-S Dd2 parasites occurs between 28-39 hpi of the IDC, with fully assembled proteasomes in schizonts remaining functional (Fig. 6B). ART-R Dd2K13^R539T^ parasites showed greater sensitivity to PfUMP1 knockdown at all stages except the late schizont stage (Fig. 6C). Notably, the ring stage displayed increased sensitivity to GlcN pulses (Fig. 6C), suggesting that the window for essential proteasome assembly in ART-R Dd2K13^R539T^ parasites is extended and occurs earlier in the IDC cycle, from late ring to mid-schizont stage (20-39 hpi).

This emphasizes a critical dependence of parasites on new 20S core particle assembly, especially in the context of the ART resistance - K13^R539T^ mutation.

## Discussion

Previous studies have mainly examined parasite responses to stress; our work offers a comprehensive characterization of parasite proteasome function and how proteasome activity varies across the pathogenic erythrocytic developmental stages of *P. falciparum.* Additionally, beyond employing the UPS in a heightened stress response, ART-R parasites have adjusted their baseline proteasome activity during ring stages, potentially as a compensatory mechanism for ART resistance.

The significant differences in proteasome activity across the IDC we observed underscore the importance of proteasomes in regulating and coordinating parasite development during the pathogenic intraerythrocytic cycle. High proteasome activity in schizonts coincides with intense DNA replication and protein turnover, processes that require maximum proteolytic activity. This observation is consistent with previous reports showing a peak in ubiquitination during the schizont stage, when high proteasome abundance and activity are required for proper schizont segregation into new daughter cells ^36,48^. Unexpectedly, ring-stage parasites maintain substantial proteasome activity despite their relatively quiescent metabolic state, possibly to manage proteotoxic stress upon erythrocyte invasion or to support early developmental remodeling ^23^.

Integrating published transcriptomic, proteomic, and metabolomic datasets with our findings, we propose that preassembled proteasomes are carried over as the parasite invades a new red blood cell, accounting for the proteasome activity observed in the ring stage. As the IDC progresses, transcription of proteasome subunit genes increases, and assembly of new proteasomes is critical for the trophozoite parasite to develop into the mature schizont form.

By conditionally depleting PfUMP1, we directly demonstrate its vital role in proteasome assembly and parasite survival. PfUMP1 knockdown caused nearly complete loss of proteasome activity and halted development at the schizont stage, confirming its conserved function as a 20S core maturation factor. Similar to its yeast and mammalian counterparts, PfUMP1 is required for proper β-subunit processing and facilitates the final steps of 20S core assembly. The critical role of PfUMP1 in *P. falciparum* viability underscores the close link between proteasome biogenesis and cell-cycle progression. Importantly, *P. falciparum* did not compensate for PfUMP1 loss and subsequent decreased proteasome activity at the transcriptional level. RNA-seq analysis showed no significant increase in proteasome subunit gene expression or other UPS components after PfUMP1 depletion, sharply contrasting with the proteasome stress responses triggered by Rpn4 in yeast or Nrf1/2 in mammals.

Our finding that *P. falciparum* does not activate canonical transcriptional feedback in response to PfUMP1 knockdown suggests that the parasite employs alternative strategies to regulate proteasome abundance and function. Although our work did not investigate the proteasome subcellular localization or post-translational modifications, both of which are known to modulate proteasome dynamics in other eukaryotes, these represent important directions for future investigation. Published phosphoproteomic studies indicate that some of the 20S subunits are phosphorylated in *P. falciparum*. Still, the stage-specificity of these modifications and their impact on proteasome assembly or catalytic activity are unknown in the parasite^49^. Similarly, the ubiquitination of proteasome subunits and other components of the UPS is documented, with the highest levels in merozoites, consistent with potential degradation of proteasome components post-invasion, accounting for the loss of activity in trophozoite forms^36^. In the mammalian system, depending on the tissue or cell type, the majority (41-74%) of cellular proteasomes exist as free 20S lacking RPs. The 19S RP is the most frequently associated 20S activator, accounting for approximately 21–57% of 20S complexes across various cells and tissues ^23,50,51^. In *P. falciparum,* the role and regulation of the 19S, PA28, and PA200 RPs have not been extensively studied except for a detailed structural characterization of *Pf*PA28. Interestingly, *Pf*PA28 is dispensable for parasite development under unstressed conditions, yet its genetic deletion increases parasite sensitivity to DHA, consistent with PA28’s canonical role as a stress response cap that facilitates ubiquitin-independent proteasome activity ^22^. Furthermore, the spatial localization of proteasome complexes is a critical determinant of proteasome activity and substrate specificity in other systems^50,51^.

Given that proteasome activity and abundance decrease during the trophozoite stage, regulated proteasome degradation may constitute an additional layer of control over proteasome homeostasis that is unknown in malaria. In eukaryotic cells, two major pathways degrade most cellular proteins: (1) the UPS, which degrades the majority of proteins, and (2) autophagy, which primarily degrades long-lived or aggregated proteins and cellular organelles through a set of Autophagy-related (ATG) proteins. In *Plasmodium* species, ATG proteins form part of a highly divergent and reduced autophagy machinery compared to other eukaryotes. Although canonical autophagy, as defined in yeast or mammals, is not fully conserved, several *Plasmodium* homologs of core ATG proteins (PfATG3, ATG5, ATG7, ATG8, ATG18, and ATG12) have been identified and appear to perform noncanonical roles in parasite survival and organelle maintenance ^52–56^. However, interactions with the proteasome have not yet been studied and warrant further investigation.

Activated ART induces widespread oxidative damage, leading to parasite death. The widely accepted mechanism of ART resistance involves mutations in the *PfK13* gene, which reduce K13-mediated hemoglobin uptake and thereby limit ART activation^7,10,41,57^. This resistance is not associated with a shift in EC_50_, as the delayed parasite clearance phenotype is linked to reduced ART sensitivity only during the early ring stage of the IDC. In isogenic parasites bearing the K13 mutation linked to ART resistance, we observed that the baseline proteasome activity was significantly elevated exclusively in the ring stage. While proteasome activity generally mirrored proteasome abundance across the IDC, ART-R parasites Dd2K13^R539T^ exhibited disproportionately elevated proteasome activity relative to abundance in ring stages. Moreover, conditional PfUMP1 knockdown revealed that ART-R parasites are markedly more sensitive to reductions in proteasome capacity than their sensitive counterparts, underscoring an increased dependency on proteasome biogenesis for survival. This strongly agrees with our previous findings that ART-R parasites (Cam3.1 R539T) were more sensitive to a proteasome inhibitor in a ring-survival assay^8^. Additionally, our data reveals that *P. falciparum* assembles the bulk of its functional proteasomes during the trophozoite–schizont transition, which are then packaged into merozoites and inherited by newly invaded rings. Blocking schizont rupture while covalently inactivating these preassembled proteasomes produces rings that form normally but cannot progress through the IDC, demonstrating that ring-stage survival depends almost entirely on proteasome pools inherited at invasion. This dependency has important implications for ART action and resistance. ART primarily targets ring-stage parasites and induces widespread protein damage and proteotoxic stress. Ring-stage survival therefore requires robust proteasome function to clear damaged proteins. Our results suggest that this capacity is predetermined by the amount of preassembled proteasome inherited at invasion.

Consequently, any perturbation that lowers the pool of assembled proteasomes entering the ring stage, such as impaired assembly due to reduced UMP1 function or direct inactivation of proteasomes renders rings unable to cope with proteotoxic stress and leads to death Though previous studies reported increased stress responses in ART-R parasites post DHA treatment/exposure, our findings demonstrate that increased proteasome activity is a baseline feature of ART-R rings, independent of acute drug stress. Further work is needed to understand how mutations in K13 increase proteasome activity. One hypothesis is that the K13-mutant early ring could be in a nutritionally deficient state, secondary to reduced hemoglobin uptake^13^, the primary source of amino acids, and consequently rely more heavily on proteasome-mediated amino acid recycling. Thus, increased proteasome activity gives ART-R parasites a survival advantage and enhanced recovery post-stress. Additionally, as K13 is also a Kelch-domain protein, it could potentially modify proteasome activity more directly through interactions with ubiquitin ligases, although canonical Nrf2-like signaling pathways are absent in *P. falciaprum*^24,27^.

ART resistance is multifactorial, involving altered stress responses, nutrient metabolism, and redox regulation, compounded by the pharmacokinetics of short-lived ART derivatives. By studying parasite proteasome biology, we gained insight into how a mutation in K13 exerts a pleomorphic effect, altering baseline proteasome dynamics in the parasite, likely enhancing parasite survival. As different resistance-conferring mutations emerge, we should study these novel mutations to determine what common aspect of parasite biology enhances survival in the face of drug pressure. We propose that the UPS and associated stress response pathways, particularly the proteasome, represent a common axis of *P. falciparum* survival under drug pressure and a promising target for overcoming ART resistance.

## Methods

### Parasite culture

*Plasmodium falciparum* parasites were cultured under standard conditions at 5% hematocrit in RPMI 1640 medium (Corning) supplemented with 24 mM sodium bicarbonate, 0.37 mM hypoxanthine, 25 mM HEPES, 10 µg/ml gentamicin and 5g/L Albumax II (Thermo Fisher Scientific). Cultures were maintained at 37°C in a gas mixture of 5% O_2_, 5% CO_2_ and 90% N_2_.

### Generation of PfUMP1-HA-glmS parasite line

The PfUMP1-HA-glmS knockdown parasite line was generated by single cross-over homologous recombination using the pSLI-HA-glmS vector^58^. A 575-bp fragment corresponding to the C-terminal region of the PfUMP1 gene (PfDd2_050032900), excluding the stop codon, was amplified from Dd2 genomic DNA using primers UMP1_F1_glmS and UMP1_R1_glmS. The fragment was inserted in frame with HA-glmS into the pSLI-HA-glmS vector (a gift from Professor R. Dzikowski) using NotI and XmaI restriction sites to generate the pSLI-PfUMP1-HA-glmS construct.

For transfection, 100µg of pSLI -PfUMP1-HA-glmS plasmid DNA was precipitated and resuspended in 400ul of Cytomix (120mM KCl, 0.15mM CaCl_2_, 2mM EGTA, 5mM MgCl_2_, 10mM K_2_HPO_4_, 25mM HEPES, pH 7.6). Parasites (Dd2 and Dd2K13^R539T^) were transfected via electroporation (0.32kV, 950µF; Bio-Rad gene-pulser) as previously described^59^. Transfected parasites were selected with 4 nM WR99210 (Jacobus Pharmaceuticals, Princeton, NJ).

Integration was confirmed by diagnostic PCR using primer pairs (a) PfUMP1_5intF/5intR and (b) PfUMP1_3intF/3intR (see Figure 2A for primer locations and Table S1 for sequences).

To select integrants, neomycin (G418; Sigma-Aldrich) was added at a final concentration of 0.4 mg/ml. Integration was confirmed by PCR as described above, and positive lines were maintained under G418 selection. From the pooled population of transfected parasites, individual clonal populations were achieved through limiting dilution and are referred to as Dd2 PfUMP1-HA-glmS and Dd2K13^R539T^ PfUMP1-HA-glmS.

### Asexual blood stage growth assay

Sorbitol-synchronized ring-stage parasites of Dd2 (WT), Dd2 PfUMP1-HA-glmS, Dd2K13^R539T^ and Dd2K13^R539T^ PfUMP1-HA-glmS were set to an initial parasitemia of 0.5 % (5% hematocrit) and cultured under standard conditions. Cultures were monitored every 48 hrs for parasitemia by flow cytometry for four replication cycles. Parasitemia was reset to 0.5% every 48 hrs to maintain parasite growth and cumulative growth was calculated. Growth curves were analyzed using MS Excel and GraphPad Prism.

### Conditional knockdown growth assay

To assess the effect of PfUMP1 knockdown, parasite growth was monitored in the glucosamine (GlcN)-treated and untreated PfUMP1-HA-glmS lines. Briefly, parasites were double synchronized with 5% sorbitol to achieve a synchrony of ± 6h in the previous cycle. Synchronized ring stage parasites of Dd2 PfUMP1-HA-glmS and Dd2K13^R539T^ PfUMP1-HA-glmS were set up at 0.5% parasitemia (5% hematocrit). Parasites were cultured in complete medium containing 0.5 mM glucosamine (GlcN) or RPMI (solvent control) for 24 hrs, after which GlcN was removed by washing twice with RPMI. Parasite growth was monitored for 3 days at 24 hrs intervals by flow cytometry and Giemsa-stained thin blood smears. Growth curves were analyzed and generated using MS Excel and GraphPad Prism.

### Stage-specific PfUMP1 knockdown assay

To determine the effect of PfUMP1 knockdown in different stages of the IDC, parasites (Dd2 PfUMP1-HA-glmS and Dd2K13^R539T^ PfUMP1-HA-glmS) were tightly synchronized.

Schizonts were purified by 70/40% Percoll-sorbitol density gradient centrifugation (4700 x g, 15 min at RT). Purified schizont stages were washed once with RPMI 1640 and allowed to reinvade fresh RBCs. After 3 hrs, residual schizonts were removed with 5% sorbitol, yielding 0–3 hpi rings. Following this, cultures were treated with 0.5 mM GlcN or RPMI control for 24 hrs at distinct developmental stages (0–3 hpi, 12–15 hpi, 27–30 hpi, and 40–43 hpi), after which GlcN was washed off and growth was monitored every 24 hrs for 3 days by flow cytometry. Growth curves were analyzed and generated using MS Excel and GraphPad Prism

### Flow cytometry

Live infected RBCs were stained for 30 min at RT in the dark with Hoechst 33342 (4 µM; Molecular Probes) and thiazole orange (100 ng/mL; Sigma-Aldrich). Parasitemia was quantified on a Cytek Aurora cytometer based on DNA (Hoechst) and RNA (thiazole orange) content. For each sample, 50,000 single cells were recorded, and doublets were excluded using FSC-H vs. FSC-W gating. Data were analyzed using SpectroFlo® software.

### Western Blot

Infected RBCs were lysed in 0.1% saponin, and parasite pellets were collected by centrifugation and washed twice in PBS. Pellets were resuspended in RIPA buffer supplemented with protease inhibitor (cOmplete™, EDTA-free; Roche), incubated on ice for 10 min, and centrifuged (13,000 × g, 15 min, 4°C). Protein concentration in the supernatant was determined using the Bio-Rad Protein Assay kit. Protein samples were then diluted in 5X SDS sample buffer, boiled at 95°C for 10 min and resolved in 4-20% polyacrylamide gel (SDS-PAGE). Proteins were transferred to 0.2 uM PVDF membrane using Invitrogen iBlot2 system and then blocked in 5 % skimmed milk/1X PBS-Tween20 at RT for 1h. Membranes were then incubated with primary antibodies (mouse anti-HA (26183, Invitrogen), rabbit anti-PfAldolase (ab207494, Abcam), mouse anti-ubiquitin (SKU VU-0101, LifeSensors), mouse anti-20S Proteasome α1/α2/α3/α5/α6/α7 (MCP231) (sc-58412, Santa Cruz) or rabbit anti-Pf proteasome ß1) overnight at 4 °C followed by incubation with HRP-conjugated secondary antibodies (Goat anti-mouse HRP or Swine anti-rabbit HRP, Dako Denmark) at RT for 1h. Immunoblots were visualized using SuperSignal West Pico PLUS chemiluminescent substrate (Thermo Fisher Scientific) and ChemiDoc MP Imaging system (Bio-Rad). Protein band intensities were quantified using Fiji (Image J).

### Analysis of Proteasome Activity by Native Gel Electrophoresis and Proteasome-Glo Assay

#### Parasite harvesting

**(a) Proteasome activity across the IDC:** Parasites were tightly synchronized. Schizonts were purified by 70/40% Percoll-sorbitol density gradient centrifugation (4700 × g, 15 min, RT), washed once with RPMI 1640, and allowed to reinvade fresh RBCs. After 3 h, residual schizonts were removed with 5% sorbitol, yielding 0–3 h post-invasion (hpi) rings. Parasites were harvested at 12–16 hpi (rings), 27–30 hpi (trophozoites), and 39–42 hpi (schizonts) by lysing infected RBCs with 0.1% saponin, followed by centrifugation and two washes in ice-cold PBS. To load equal total protein required vastly different volumes of parasite culture, given the size and the small amount of cellular material in a ring-stage parasite.
**(b) Proteasome activity upon PfUMP1 knockdown:** Double-sorbitol synchronized ring-stage Dd2 PfUMP1-HA-glmS parasites were cultured with or without 0.5 mM glucosamine (GlcN) for 24 h and harvested at the early schizont stage (34–39 hpi). As controls, late trophozoite-stage Dd2 and Dd2 PfUMP1-HA-glmS parasites were treated with either 1 µM TDI8304 (a Pf-specific proteasome inhibitor) or DMSO (vehicle control) for 4 h at 37°C, then harvested at the early schizont stage. Parasite pellets were collected as described above.

#### Preparation of parasite extracts

Parasite pellets were resuspended in lysis buffer (50 mM Tris, 2 mM DTT, 5 mM MgCl, 10% glycerol, 2 mM ATP, 0.05% digitonin, pH 7.5). To preserve the integrity of the 26S proteasome complex, a mild extraction protocol was used: cells were incubated on ice for 30 min with gentle vortexing every 10 min. Lysates were clarified by centrifugation (13,000 × g, 15 min, 4°C), and protein concentrations were determined by BCA assay (Pierce).

#### Native gel electrophoresis and in-gel proteasome activity assay

For native PAGE, 40 µg of parasite lysate was mixed with native loading buffer (0.05% bromophenol blue, 43.5% glycerol, 250 mM Tris, pH 7.5) and resolved on a NuPAGE 3–8% Tris-acetate gel (NOVEX Life Technologies). Electrophoresis was performed in TBE buffer supplemented with 0.4 mM ATP, 2 mM MgCl, and 0.5 mM DTT at 150 V for 3 hrs at 4°C. Following electrophoresis, gels were incubated in developing buffer (50 mM Tris, pH 7.5, 10 mM MgCl, 1 mM ATP, 1 mM DTT) containing 50 µM Suc-LLVY-AMC for 30 min at 37°C in the dark. Fluorescent signals were detected under UV light using the Bio-Rad ChemiDoc MP imaging system. Band intensities were quantified by densitometry using Bio-Rad Image Lab software and normalized to WT Dd2 (set as 100% activity).

#### Proteasome-Glo assay

For the Proteasome-Glo assay, 1 µg of parasite lysate in 50 µL was mixed with an equal volume of Proteasome-Glo (chymotrypsin-like) reagent (Promega, Madison, WI), following the manufacturer’s instructions. Luminescence was measured with a microplate reader at 60 min, corresponding to the peak signal.

### Quantitative RNA-seq

For RNA seq experiments, tightly synchronized cultures of Dd2 and PfUMP1-HA-glmS were obtained by Percoll-sorbitol synchronization, and parasites were harvested at 31-34 hpi. For knockdown of PfUmp1, tightly synchronized PfUMP1-HA-glmS parasites were treated with 0.5 mM GlcN at 7-10 hpi for 24 hrs and harvested at 31-34 hpi. Parasites were harvested by saponin lysis and TRIzol extraction method was used for total RNA isolation. Following RNA isolation, total RNA integrity was checked using a 2100 Bioanalyzer (Agilent Technologies, Santa Clara, CA). RNA concentrations were measured using the NanoDrop system (Thermo Fisher Scientific, Inc., Waltham, MA). The Genomics Core Laboratory performed preparation of RNA sample library and RNA-seq at Weill Cornell Medicine. The sample was processed with the Agilent high-throughput sample preparation Bravo B system, automated with the Illumina TruSeq Stranded mRNA Sample Library Preparation kit (Illumina, San Diego, CA), according to the manufacturer’s instructions. The normalized cDNA libraries were pooled and sequenced on an Illumina NovaSeq 6000 sequencer with paired-end; 100-cycle reads. The raw sequencing reads in BCL format were processed through bcl2fastq 2.19 (Illumina) for FASTQ conversion and demultiplexing. After trimming the adaptors with cutadapt (version1.18) (https://cutadapt.readthedocs.io/en/v1.18/), RNA reads were aligned and mapped to the Plasmodium falciparum Dd2 reference genome (https://plasmodb.org/plasmo/app/downloads/Current_Release/PfalciparumDd2/fasta/data/) by STAR (Version2.5.2) (https://github.com/alexdobin/STAR). The abundance of transcripts was measured with salmon v1.9.0 (https://github.com/COMBINE-lab/salmon) in Transcripts Per Million (TPM). Raw read counts per gene were extracted using HTSeq-count v0.11.2. Gene expression profiles were constructed for differential expression, cluster, and principle component analyses with the DESeq2 package (https://bioconductor.org/packages/release/bioc/html/DESeq2.html). For differential expression analysis, pairwise comparisons between two or more groups using parametric tests where read-counts follow a negative binomial distribution with a gene-specific dispersion parameter. Corrected p-values were calculated based on the Benjamini-Hochberg method to adjust for multiple testing.

### Pf ß1 proteasome subunit protein expression, purification and antibody generation

The coding sequence of Pf ß1 proteasome subunit (PF3D7_0931800) was codon-optimized and cloned into the pRFSDuet-1 expression vector using BamHI and HindIII restriction sites. The full-length Pfß1 protein was expressed in *E.coli* BL21 cells as an N-terminal His-Sumo-tagged protein (His-Sumo Pfß1) in. Recombinant His-Sumo Pfß1was purified using Ni-NTA affinity chromatography following the manufacturer’s protocol (HisPur™ Ni-NTA Spin Columns, Thermo Fisher Scientific, Cat#88226). The His-SUMO tag was cleaved with SUMO protease, and the untagged Pfβ1 protein was further purified by Ni-NTA affinity chromatography.

Polyclonal antibodies against purified Pfβ1 were generated in rabbits by Pocono Rabbit Laboratory using their 28-day Mighty Quick protocol. Briefly, a rabbit was immunized with 200 µg Pfβ1 protein formulated with Mighty Quick Immune Stimulator in complete Freund’s adjuvant, followed by booster injections of 100 µg Pfβ1 protein with Mighty Quick Immune Stimulator in incomplete Freund’s adjuvant at weeks 1 and 2. On day 28, the rabbit was exsanguinated, and serum was collected. Pfβ1-specific antibodies were enriched from serum by affinity purification using recombinant His-SUMO–Pfβ1 protein. Purified antibody was supplemented with 0.01% sodium azide and50% glycerol for storage at −80°C.

## References

1. WHO. World malaria report 2023. (Geneva: World Health Organization, 2023).

2 Hamilton, W. L. et al. Evolution and expansion of multidrug-resistant malaria in southeast Asia: a genomic epidemiology study. Lancet Infect Dis 19, 943–951 (2019). 10.1016/S1473-3099(19)30392-5

3 Balikagala, B. et al. Evidence of Artemisinin-Resistant Malaria in Africa. N Engl J Med 385, 1163–1171 (2021). 10.1056/NEJMoa2101746

4 Uwimana, A. et al. Emergence and clonal expansion of in vitro artemisinin-resistant Plasmodium falciparum kelch13 R561H mutant parasites in Rwanda. Nat Med 26, 1602–1608 (2020). 10.1038/s41591-020-1005-2

5 Woodrow, C. J. & White, N. J. The clinical impact of artemisinin resistance in Southeast Asia and the potential for future spread. FEMS Microbiol Rev 41, 34–48 (2017). 10.1093/femsre/fuw037

6 Dogovski, C. et al. Targeting the cell stress response of Plasmodium falciparum to overcome artemisinin resistance. PLoS Biol 13, e1002132 (2015). 10.1371/journal.pbio.1002132

7 Mok, S. et al. Artemisinin-resistant K13 mutations rewire Plasmodium falciparum’s intra-erythrocytic metabolic program to enhance survival. Nat Commun 12, 530 (2021). 10.1038/s41467-020-20805-w

8 Zhan, W. et al. Dual-pharmacophore artezomibs hijack the Plasmodium ubiquitin-proteasome system to kill malaria parasites while overcoming drug resistance. Cell Chem Biol 30, 457–469 e411 (2023). 10.1016/j.chembiol.2023.04.006

9 Straimer, J. et al. Drug resistance. K13-propeller mutations confer artemisinin resistance in Plasmodium falciparum clinical isolates. Science 347, 428–431 (2015). 10.1126/science.1260867

10 Bridgford, J. L. et al. Artemisinin kills malaria parasites by damaging proteins and inhibiting the proteasome. Nat Commun 9, 3801 (2018). 10.1038/s41467-018-06221-1

11 Meshnick, S. R. Artemisinin: mechanisms of action, resistance and toxicity. Int J Parasitol 32, 1655–1660 (2002). 10.1016/s0020-7519(02)00194-7

12 Yang, Y. Z., Little, B. & Meshnick, S. R. Alkylation of proteins by artemisinin. Effects of heme, pH, and drug structure. Biochem Pharmacol 48, 569–573 (1994). 10.1016/0006-2952(94)90287-9

13 Birnbaum, J. et al. A Kelch13-defined endocytosis pathway mediates artemisinin resistance in malaria parasites. Science 367, 51–59 (2020). 10.1126/science.aax4735

14 Talman, A. M., Clain, J., Duval, R., Menard, R. & Ariey, F. Artemisinin Bioactivity and Resistance in Malaria Parasites. Trends Parasitol 35, 953–963 (2019). 10.1016/j.pt.2019.09.005

15 Witkowski, B. et al. Novel phenotypic assays for the detection of artemisinin-resistant Plasmodium falciparum malaria in Cambodia: in-vitro and ex-vivo drug-response studies. Lancet Infect Dis 13, 1043–1049 (2013). 10.1016/S1473-3099(13)70252-4

16 Rocamora, F. et al. Oxidative stress and protein damage responses mediate artemisinin resistance in malaria parasites. PLoS Pathog 14, e1006930 (2018). 10.1371/journal.ppat.1006930

17 Hsu, H. C. et al. Structures revealing mechanisms of resistance and collateral sensitivity of Plasmodium falciparum to proteasome inhibitors. Nat Commun 14, 8302 (2023). 10.1038/s41467-023-44077-2

18 Kirkman, L. A. et al. Antimalarial proteasome inhibitor reveals collateral sensitivity from intersubunit interactions and fitness cost of resistance. Proc Natl Acad Sci U S A 115, E6863–E6870 (2018). 10.1073/pnas.1806109115

19 Lawong, A. et al. Identification of potent and reversible piperidine carboxamides that are species-selective orally active proteasome inhibitors to treat malaria. Cell Chem Biol (2024). 10.1016/j.chembiol.2024.07.001

20 Li, H. et al. Structure- and function-based design of Plasmodium-selective proteasome inhibitors. Nature 530, 233–236 (2016). 10.1038/nature16936

21 Tanaka, K. The proteasome: overview of structure and functions. Proc Jpn Acad Ser B Phys Biol Sci 85, 12–36 (2009). 10.2183/pjab.85.12

22 Xie, S. C. et al. The structure of the PA28-20S proteasome complex from Plasmodium falciparum and implications for proteostasis. Nat Microbiol 4, 1990–2000 (2019). 10.1038/s41564-019-0524-4

23 Livneh, I., Cohen-Kaplan, V., Cohen-Rosenzweig, C., Avni, N. & Ciechanover, A. The life cycle of the 26S proteasome: from birth, through regulation and function, and onto its death. Cell Res 26, 869–885 (2016). 10.1038/cr.2016.86

24 Jang, J., Wang, Y., Kim, H. S., Lalli, M. A. & Kosik, K. S. Nrf2, a regulator of the proteasome, controls self-renewal and pluripotency in human embryonic stem cells. Stem Cells 32, 2616–2625 (2014). 10.1002/stem.1764

25 Kapeta, S., Chondrogianni, N. & Gonos, E. S. Nuclear erythroid factor 2-mediated proteasome activation delays senescence in human fibroblasts. J Biol Chem 285, 8171–8184 (2010). 10.1074/jbc.M109.031575

26 Mannhaupt, G., Schnall, R., Karpov, V., Vetter, I. & Feldmann, H. Rpn4p acts as a transcription factor by binding to PACE, a nonamer box found upstream of 26S proteasomal and other genes in yeast. FEBS Lett 450, 27–34 (1999). 10.1016/s0014-5793(99)00467-6

27 Pickering, A. M., Linder, R. A., Zhang, H., Forman, H. J. & Davies, K. J. A. Nrf2-dependent induction of proteasome and Pa28alphabeta regulator are required for adaptation to oxidative stress. J Biol Chem 287, 10021–10031 (2012). 10.1074/jbc.M111.277145

28 Radhakrishnan, S. K. et al. Transcription factor Nrf1 mediates the proteasome recovery pathway after proteasome inhibition in mammalian cells. Mol Cell 38, 17–28 (2010). 10.1016/j.molcel.2010.02.029

29 Xie, Y. & Varshavsky, A. RPN4 is a ligand, substrate, and transcriptional regulator of the 26S proteasome: a negative feedback circuit. Proc Natl Acad Sci U S A 98, 3056–3061 (2001). 10.1073/pnas.071022298

30 Kunjappu, M. J. & Hochstrasser, M. Assembly of the 20S proteasome. Biochim Biophys Acta 1843, 2–12 (2014). 10.1016/j.bbamcr.2013.03.008

31 Murata, S., Yashiroda, H. & Tanaka, K. Molecular mechanisms of proteasome assembly. Nat Rev Mol Cell Biol 10, 104–115 (2009). 10.1038/nrm2630

32 Ramos, P. C., Hockendorff, J., Johnson, E. S., Varshavsky, A. & Dohmen, R. J. Ump1p is required for proper maturation of the 20S proteasome and becomes its substrate upon completion of the assembly. Cell 92, 489–499 (1998). 10.1016/s0092-8674(00)80942-3

33 Zhan, W. et al. Development of a Highly Selective Plasmodium falciparum Proteasome Inhibitor with Anti-malaria Activity in Humanized Mice. Angew Chem Int Ed Engl 60, 9279–9283 (2021). 10.1002/anie.202015845

34. Rosenthal, M. R., Vijayrajratnam, S., Firestone, T. M. & Ng, C. L. Enhanced cell stress response and protein degradation capacity underlie artemisinin resistance in Plasmodium falciparum. mSphere 9, e0037124 (2024). 10.1128/msphere.00371-24

35 Kisselev, A. F. & Goldberg, A. L. Monitoring activity and inhibition of 26S proteasomes with fluorogenic peptide substrates. Methods Enzymol 398, 364–378 (2005). 10.1016/S0076-6879(05)98030-0

36 Green, J. L. et al. Ubiquitin activation is essential for schizont maturation in Plasmodium falciparum blood-stage development. PLoS Pathog 16, e1008640 (2020). 10.1371/journal.ppat.1008640

37 Moravec, R. A. et al. Cell-based bioluminescent assays for all three proteasome activities in a homogeneous format. Anal Biochem 387, 294–302 (2009). 10.1016/j.ab.2009.01.016

38 Cowman, A. F. & Crabb, B. S. Invasion of red blood cells by malaria parasites. Cell 124, 755–766 (2006). 10.1016/j.cell.2006.02.006

39 Singh, S. & Chitnis, C. E. Signalling mechanisms involved in apical organelle discharge during host cell invasion by apicomplexan parasites. Microbes Infect 14, 820–824 (2012). 10.1016/j.micinf.2012.05.007

40 Taylor, H. M. et al. The malaria parasite cyclic GMP-dependent protein kinase plays a central role in blood-stage schizogony. Eukaryot Cell 9, 37–45 (2010). 10.1128/EC.00186-09

41 Birnbaum, J. et al. A genetic system to study Plasmodium falciparum protein function. Nat Methods 14, 450–456 (2017). 10.1038/nmeth.4223

42 Marshall, R. S. & Vierstra, R. D. Dynamic Regulation of the 26S Proteasome: From Synthesis to Degradation. Front Mol Biosci 6, 40 (2019). 10.3389/fmolb.2019.00040

43 Shirozu, R., Yashiroda, H. & Murata, S. Identification of minimum Rpn4-responsive elements in genes related to proteasome functions. FEBS Lett 589, 933–940 (2015). 10.1016/j.febslet.2015.02.025

44 Koizumi, S., Hamazaki, J. & Murata, S. Transcriptional regulation of the 26S proteasome by Nrf1. Proc Jpn Acad Ser B Phys Biol Sci 94, 325–336 (2018). 10.2183/pjab.94.021

45 Yabuta, Y. et al. Involvement of Arabidopsis NAC transcription factor in the regulation of 20S and 26S proteasomes. Plant Sci 181, 421–427 (2011). 10.1016/j.plantsci.2011.07.001

46 Iengar, P. & Joshi, N. V. Identification of putative regulatory motifs in the upstream regions of co-expressed functional groups of genes in Plasmodium falciparum. BMC Genomics 10, 18 (2009). 10.1186/1471-2164-10-18

47 Stokes, B. H. et al. Covalent Plasmodium falciparum-selective proteasome inhibitors exhibit a low propensity for generating resistance in vitro and synergize with multiple antimalarial agents. PLoS Pathog 15, e1007722 (2019). 10.1371/journal.ppat.1007722

48 Ponts, N. et al. Unraveling the ubiquitome of the human malaria parasite. J Biol Chem 286, 40320–40330 (2011). 10.1074/jbc.M111.238790

49 Solyakov, L. et al. Global kinomic and phospho-proteomic analyses of the human malaria parasite Plasmodium falciparum. Nat Commun 2, 565 (2011). 10.1038/ncomms1558

50 Morozov, A. V. & Karpov, V. L. Biological consequences of structural and functional proteasome diversity. Heliyon 4, e00894 (2018). 10.1016/j.heliyon.2018.e00894

51 Fabre, B. et al. Subcellular distribution and dynamics of active proteasome complexes unraveled by a workflow combining in vivo complex cross-linking and quantitative proteomics. Mol Cell Proteomics 12, 687–699 (2013). 10.1074/mcp.M112.023317

52 Walker, D. M. et al. Plasmodium falciparum erythrocytic stage parasites require the putative autophagy protein PfAtg7 for normal growth. PLoS One 8, e67047 (2013). 10.1371/journal.pone.0067047

53 Kitamura, K. et al. Autophagy-related Atg8 localizes to the apicoplast of the human malaria parasite Plasmodium falciparum. PLoS One 7, e42977 (2012). 10.1371/journal.pone.0042977

54 Tomlins, A. M. et al. Plasmodium falciparum ATG8 implicated in both autophagy and apicoplast formation. Autophagy 9, 1540–1552 (2013). 10.4161/auto.25832

55 Bansal, P., Tripathi, A., Thakur, V., Mohmmed, A. & Sharma, P. Autophagy-Related Protein ATG18 Regulates Apicoplast Biogenesis in Apicomplexan Parasites. mBio 8 (2017). 10.1128/mBio.01468-17

56 Agrawal, P., Manjithaya, R. & Surolia, N. Autophagy-related protein PfATG18 participates in food vacuole dynamics and autophagy-like pathway in Plasmodium falciparum. Mol Microbiol 113, 766–782 (2020). 10.1111/mmi.14441

57 Ismail, H. M. et al. Artemisinin activity-based probes identify multiple molecular targets within the asexual stage of the malaria parasites Plasmodium falciparum 3D7. Proc Natl Acad Sci U S A 113, 2080–2085 (2016). 10.1073/pnas.1600459113

58 Prommana, P. et al. Inducible knockdown of Plasmodium gene expression using the glmS ribozyme. PLoS One 8, e73783 (2013). 10.1371/journal.pone.0073783

59 Crabb, B. S. et al. Transfection of the human malaria parasite Plasmodium falciparum. Methods Mol Biol 270, 263–276 (2004). 10.1385/1-59259-793-9:263

60 Kucharski, M. et al. A comprehensive RNA handling and transcriptomics guide for high-throughput processing of Plasmodium blood-stage samples. Malar J 19, 363 (2020). 10.1186/s12936-020-03436-w

61 Chappell, L. et al. Refining the transcriptome of the human malaria parasite Plasmodium falciparum using amplification-free RNA-seq. BMC Genomics 21, 395 (2020). 10.1186/s12864-020-06787-5

