## Supplemental tables and figures for "Stage-specific regulation of the *Plasmodium falciparum* proteasome activity reveals adaptive rewiring in artemisinin resistance"

**Supplementary Figures:

Supplementary Fig. 1**

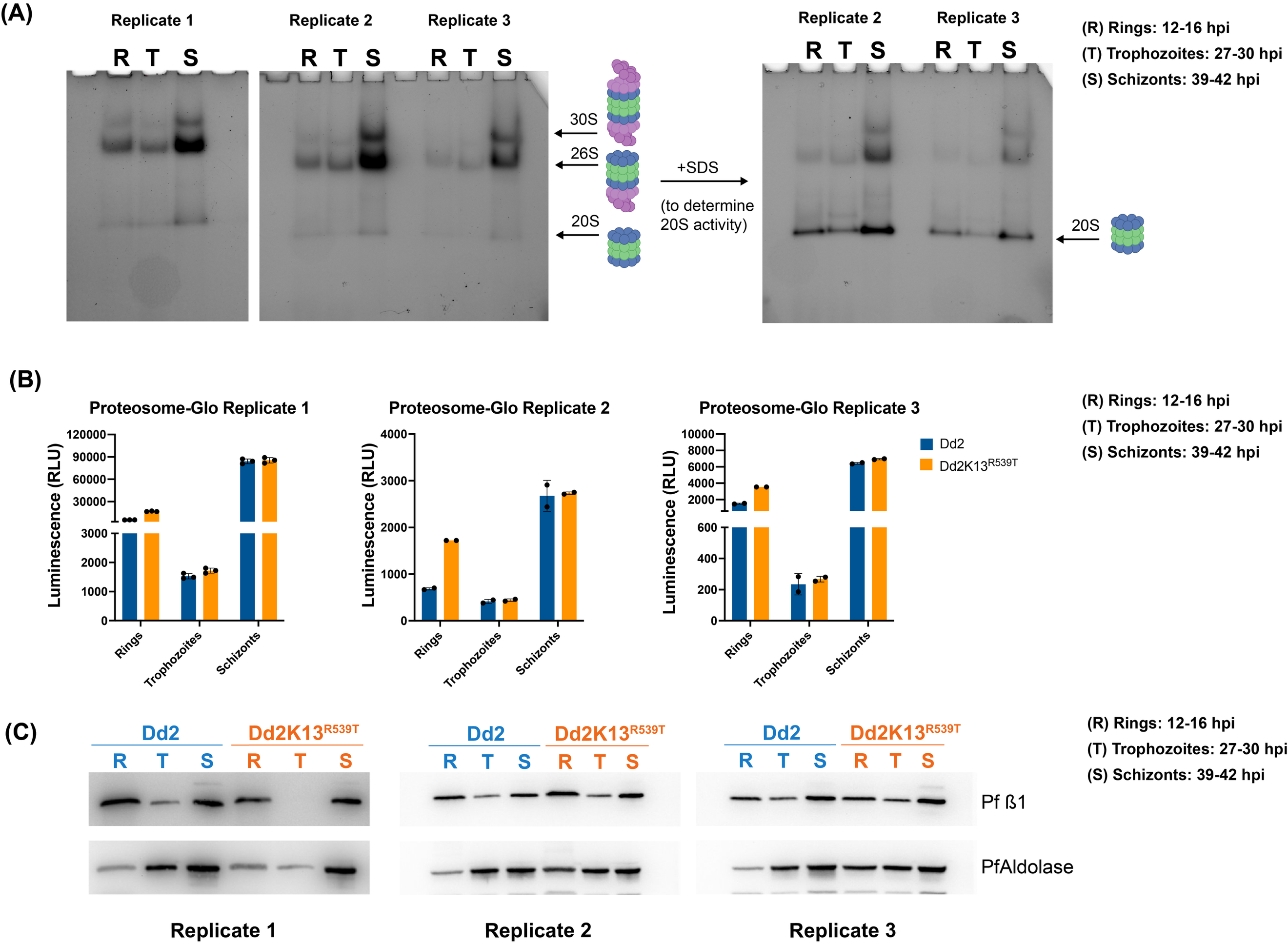

(A) Biological replicates of in-gel proteasome activity assay using SucLLVY-AMC (detects chymotrypsin activity exhibited by the ß5 subunit of the proteasome) as substrate. Extracts from Dd2 parasites (rings, trophozoites, and schizonts) were separated by native gel and peptidase activity from the proteasome was visualized using SucLLVY-AMC in gel activity assay in the absence (left) or presence (right) of 0.02% SDS (which opens and activates the 20S proteasome to visualize free 20S core particle).
(B) Biological replicates of Proteasome-Glo^TM^ assay on rings, trophozoites and schizont stages of Dd2 and Dd2K13^R539T^ parasites. Cellular lysates were incubated with Suc-LLVY-AMC for 10 min at 37°C, and fluorescence intensity was measured.
(C) Biological replicates of western blot analysis of Pf proteasome ß1 subunit expression levels across the IDC (rings, trophozoites, and schizonts) of Dd2 and Dd2K13^R539T^ parasites. PfAldolase served as a loading control.

**Supplementary Fig. 2**

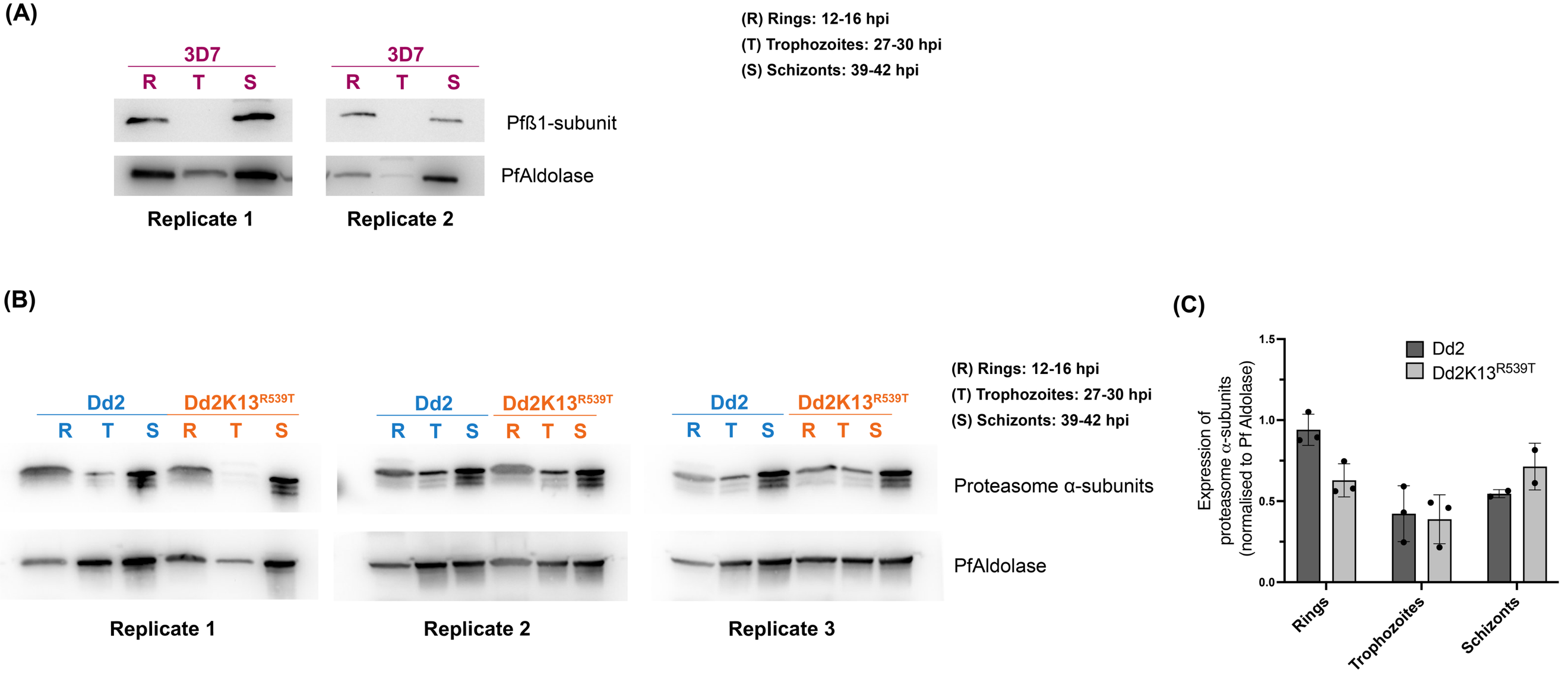

(A) Biological replicates of western blot analysis of *Pf* proteasome ß1 subunit expression levels across the IDC (rings, trophozoites, and schizonts) of 3D7 parasites. PfAldolase served as a loading control.
(B) Biological replicates of western blot analysis of proteasome α-subunits expression levels across the IDC (rings, trophozoites, and schizonts) of Dd2 and Dd2K13^R539T^ parasites. PfAldolase served as a loading control.
(C) Western blot quantification of proteasome α-subunits expression from (B)

**Supplementary Fig. 3**

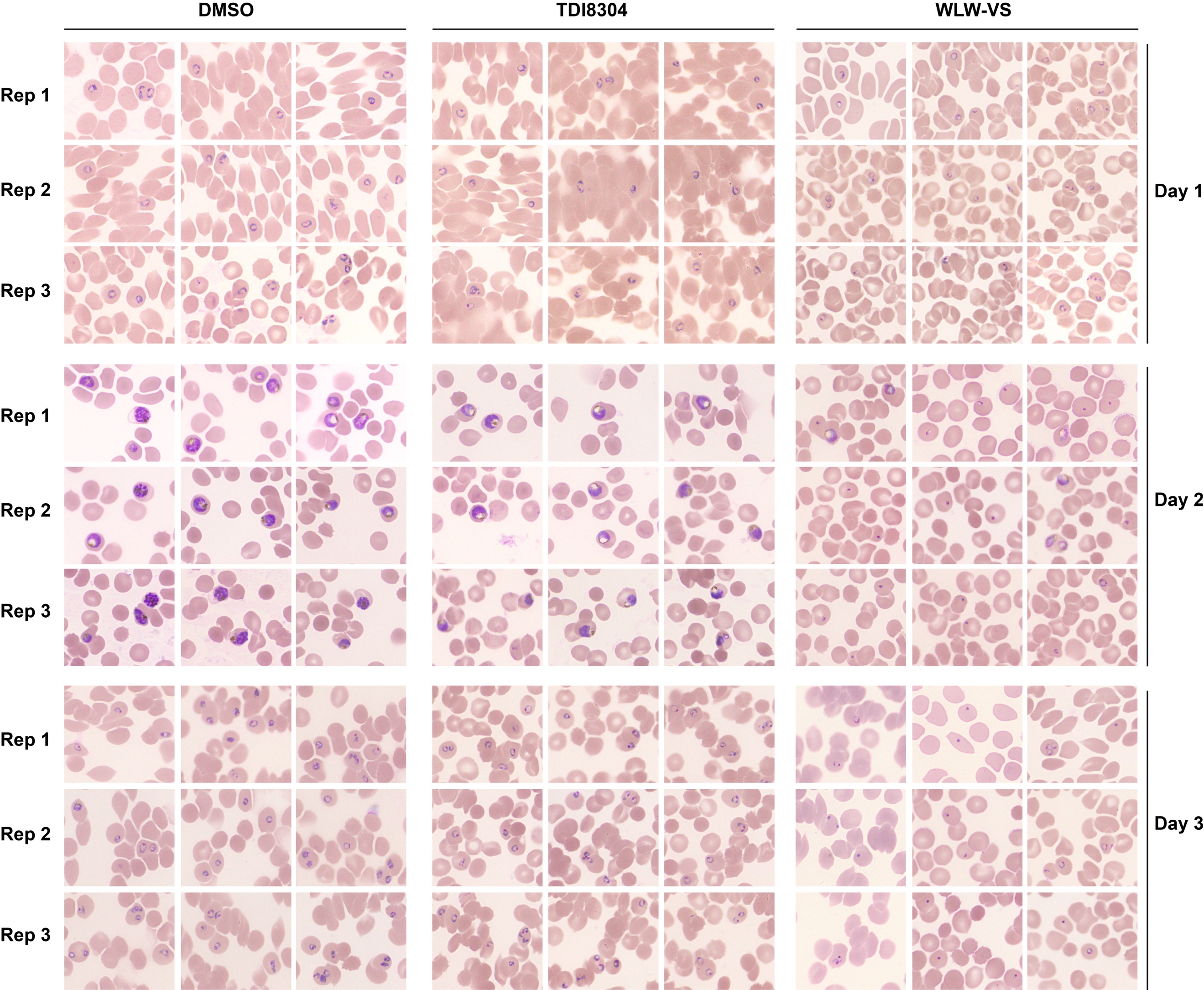

Biological replicates of Fig. 2. Giemsa-stained smears of DMSO/TDI8304/WLW-VS treated parasites on days 1-3 post inhibitor treatment. Briefly, tightly synchronized 44–46 hpi segmented schizonts of ART-S Dd2 parasites were treated with 2.5 μM compound 1 for 5 hours^40^. Following compound 1 treatment, parasites were exposed to the covalent proteasome inhibitor WLW-VS (1 μM, 1 h) to irreversibly inhibit assembled proteasomes. Controls included DMSO and the reversible proteasome inhibitor TDI-8304 (1 μM). Cultures were then washed free of inhibitors and returned to growth, and growth was monitored for 3 days. Stars represent pyknotic parasites.

**Supplementary Fig. 4**

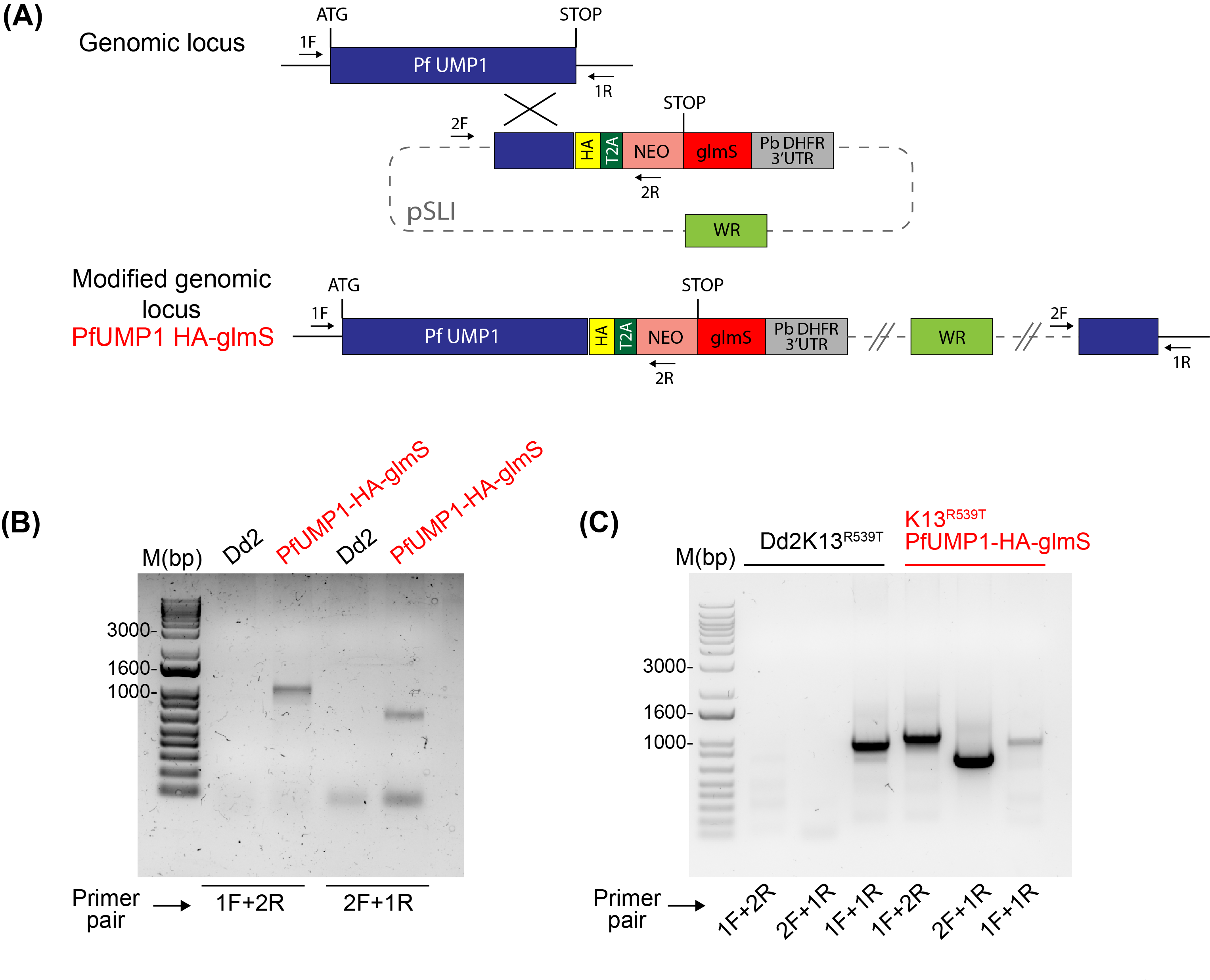

(A) Schematic of the selection-linked integration-based strategy used to generate PfUMP1-HA-glmS parasites by single-crossover recombination. Arrows denote diagnostic PCR primers. T2A, skip peptide; Neo, neomycin-resistance gene; glmS, glmS ribozyme.
(B) Representative PCR of the transfected PfUMP1-HA-glmS parasites confirming the integration of the HA-glmS sequence to the 3’end of PfUMP1. Genomic DNA from parental (Dd2) and transfected PfUMP1-HA-glmS parasites was used as the template.
(C) Representative PCR of the transfected K13^R539T^PfUMP1-HA-glmS parasites confirming the integration of the HA-glmS sequence to the 3’end of PfUMP1. Genomic DNA from parental (Dd2K13^R539T^) and transfected PfUMP1-HA-glmS parasites was used as the template.

**Supplementary Fig. 5**

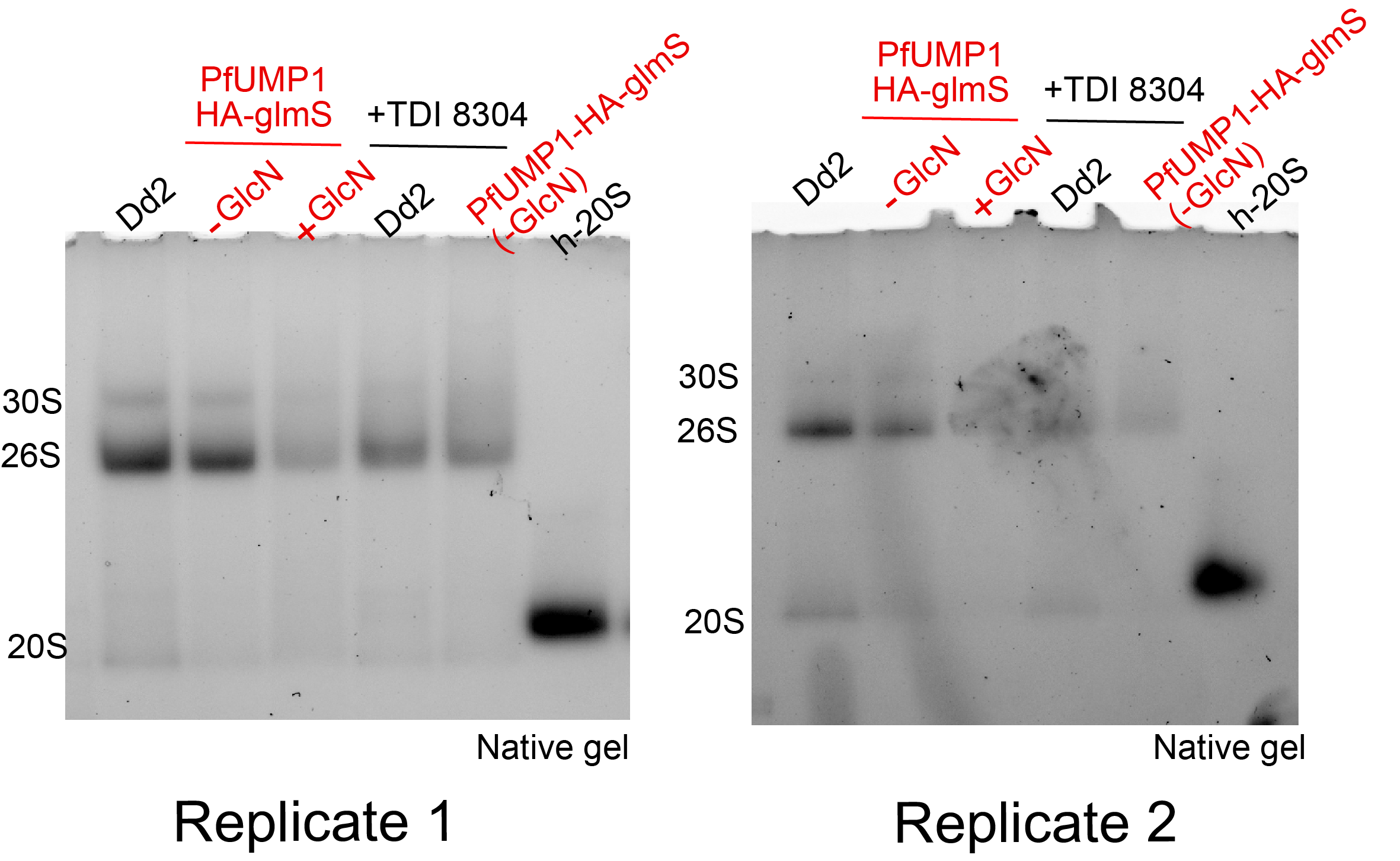

Biological replicates of in-gel proteasome activity assay using SucLLVY-AMC (detects chymotrypsin activity exhibited by the ß5 subunit of the proteasome) as substrate. Cell extracts from 34-39 hpi Dd2 and PfUMP1-HA-glmS parasites, treated with or without GlcN for 24h, were separated by native gel and peptidase activity from the proteasome was visualized using SucLLVY-AMC. Analysis of proteasome activity of Dd2 and PfUMP1-HA-glmS parasites treated with TDI-8304, a selective Plasmodium proteasome inhibitor served as a control.

**Supplementary Fig. 6**

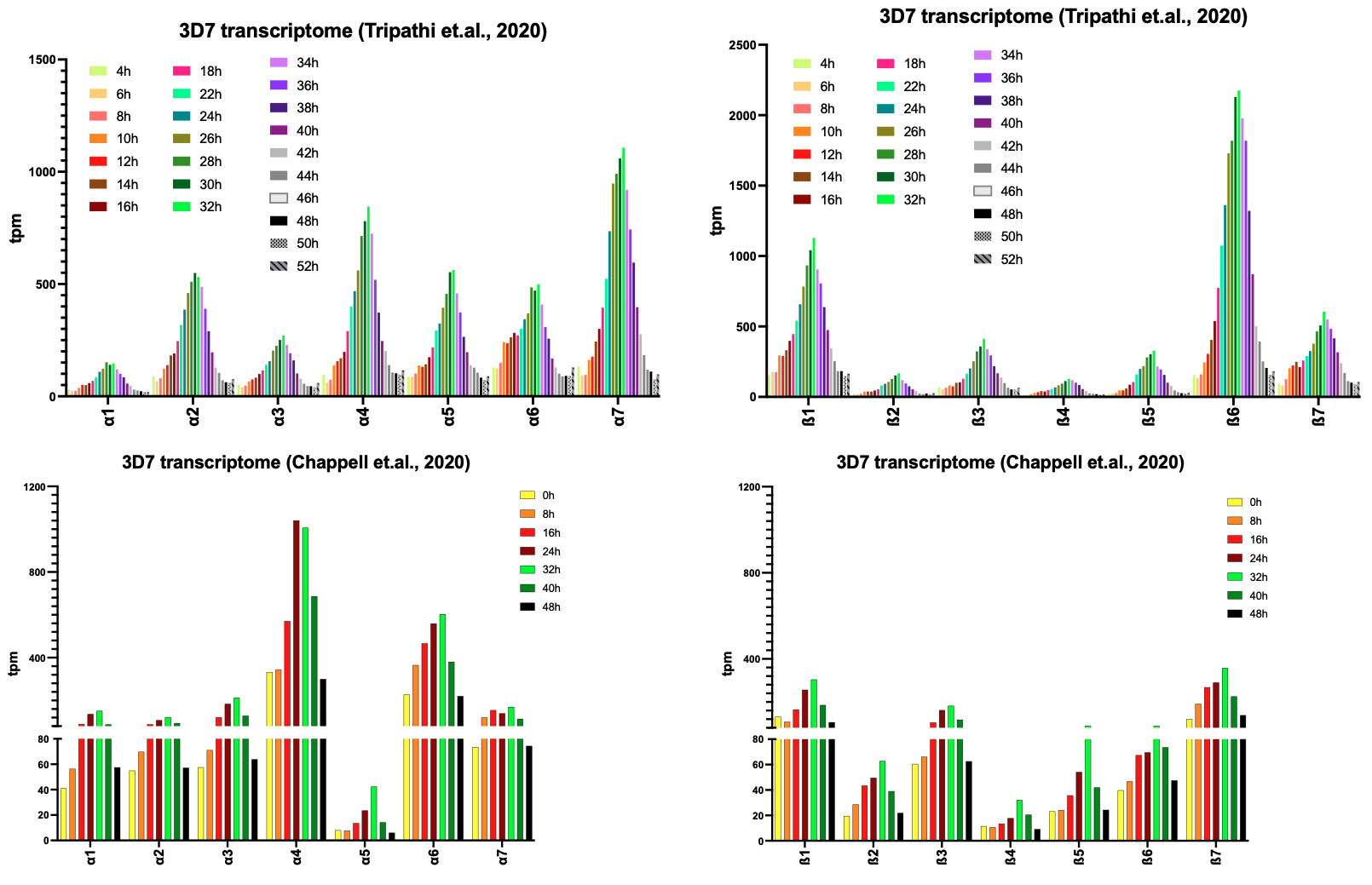

Transcriptome analysis of *Pf* proteasome α and ß subunits across the IDC from two different whole-transcriptome data sets^60,61^
